# Tuning of self-renewing capacity in the *Arabidopsis* stomatal lineage by hormonal and nutrition regulation of asymmetric cell divisions

**DOI:** 10.1101/2020.09.16.300830

**Authors:** Yan Gong, Julien Alassimone, Rachel Varnau, Nidhi Sharma, Lily S. Cheung, Dominique C. Bergmann

## Abstract

Asymmetric and self-renewing divisions build and pattern tissues. In the *Arabidopsis thaliana* stomatal lineage, asymmetric cell divisions, guided by polarly localized cortical proteins, generate the majority of cells on the leaf surface. These divisions can be fine-tuned by systemic and environmental signals to modify tissue development, but the molecular mechanisms by which plants incorporate such cues to regulate asymmetric divisions are largely unknown. In a screen for modulators of cell polarity and asymmetric divisions, we identified a mutation in *CONSTITIUTIVE TRIPLE RESPONSE 1*, a negative regulator of ethylene signaling. We subsequently revealed antagonistic impacts of ethylene and glucose signaling on the self-renewing capacity of stomatal lineage stem cells. Quantitative analysis of the impacts of these signaling systems on cell polarity and fate dynamics showed that developmental information may be encoded in both the spatial and temporal asymmetries of polarity proteins. Taken together, our results provide a framework for a mechanistic understanding of how systemic information such as nutritional status and environmental factors tune stem cell behavior in the stomatal lineage, ultimately enabling optimization of leaf size and cell-type composition.

## INTRODUCTION

The cellular composition of a tissue defines its structure and functions. By modulating their division behaviors, tissue-embedded stem cells control cell fate and distribution (Losick et al., 2011; Motohashi & Asakura, 2014), often using symmetric cell divisions (SCDs) to renew stem-cell capacity, and asymmetric cell divisions (ACDs) to diversify daughter cells’ fates (De Smet & Beeckman, 2011; Morrison & Kimble, 2006). This ability to tune the relative proportion of ACDs and SCDs can provide plasticity to regulate tissue size and organ composition in response to changes in the external and internal environment. For example, by shifting the ratio of ACDs to SCDs, intestinal stem cells in *Drosophila melanogaster* can resize the intestine in response to food availability (O’Brien et al., 2011), and in rats, brains cells are able to replenish differentiated neurons during stroke recovery (Zhang et al., 2004).

Cell polarity, the restricted localization of proteins, organelles, and activities to one region of the cell, is often linked to ACDs. During ACDs, cell polarity can precede and dictate division orientation, thereby affecting not only daughter cell size but cell fate through positional and differential inheritance of specific materials (Knoblich, 2001; Muroyama & Bergmann, 2019). After ACDs, cell polarity can also play important roles in determining the subsequent division type. If a cell undergoes successive rounds of ACDs, polarity must either be maintained or regenerated at each ACD. When the degree of polarity is not sufficient to ensure differential segregation of proteins to one daughter, it can trigger a developmental switch from ACDs to SCDs (and subsequent differentiation), as was demonstrated for PAR proteins in *Caenorhabditis elegans* embryo development (Hubatsch et al., 2019).

The stomatal lineage in the epidermis of plant leaves is an excellent model to study the interplay of cell polarity and cell division behaviors with developmental plasticity and physiological adaptation. In *Arabidopsis*, stomatal lineage stem cells give rise to different ratios of two essential cell types, stomatal guard cells (GCs) and pavement cells, by modulating the frequency of ACDs and SCDs (Figure 1A). At any given time during development, stomatal lineages at different developmental stages can be found distributed across the surface of a leaf. These lineages are initiated by ACDs which produce meristemoids and stomatal lineage ground cells (SLGCs), and successive ACDs in these daughter cells replenish self-renewing cells. Terminal differentiation coincides with the SCD, and subsequent differentiation, of a guard mother cell (GMC) into guard cells. By regulating the balance of differentiation and self-renewal (approximated by the SCD/ACD ratio) of the stomatal lineage cells, plants can change the composition, size, and patterning of the leaf epidermis (Bergmann & Sack, 2007; Vaten et al., 2018), which further determines the leaf organ size (Gonzalez et al., 2012; Vaseva et al., 2018). This process is highly flexible and responsive to many intrinsic and extrinsic factors (Engineer et al., 2014; Lau et al., 2018; Lee et al., 2017; Schroeder et al., 2001). For example, a recent analysis of cytokinin hormone signaling showed that regulating the ability of SLGCs to undergo ACDs (also known as spacing divisions) contributes to developmental flexibility (Vaten et al., 2018). However, a mechanistic understanding of how these intrinsic and extrinsic factors influence cell fate and division behaviors is lacking.

**Figure 1.**
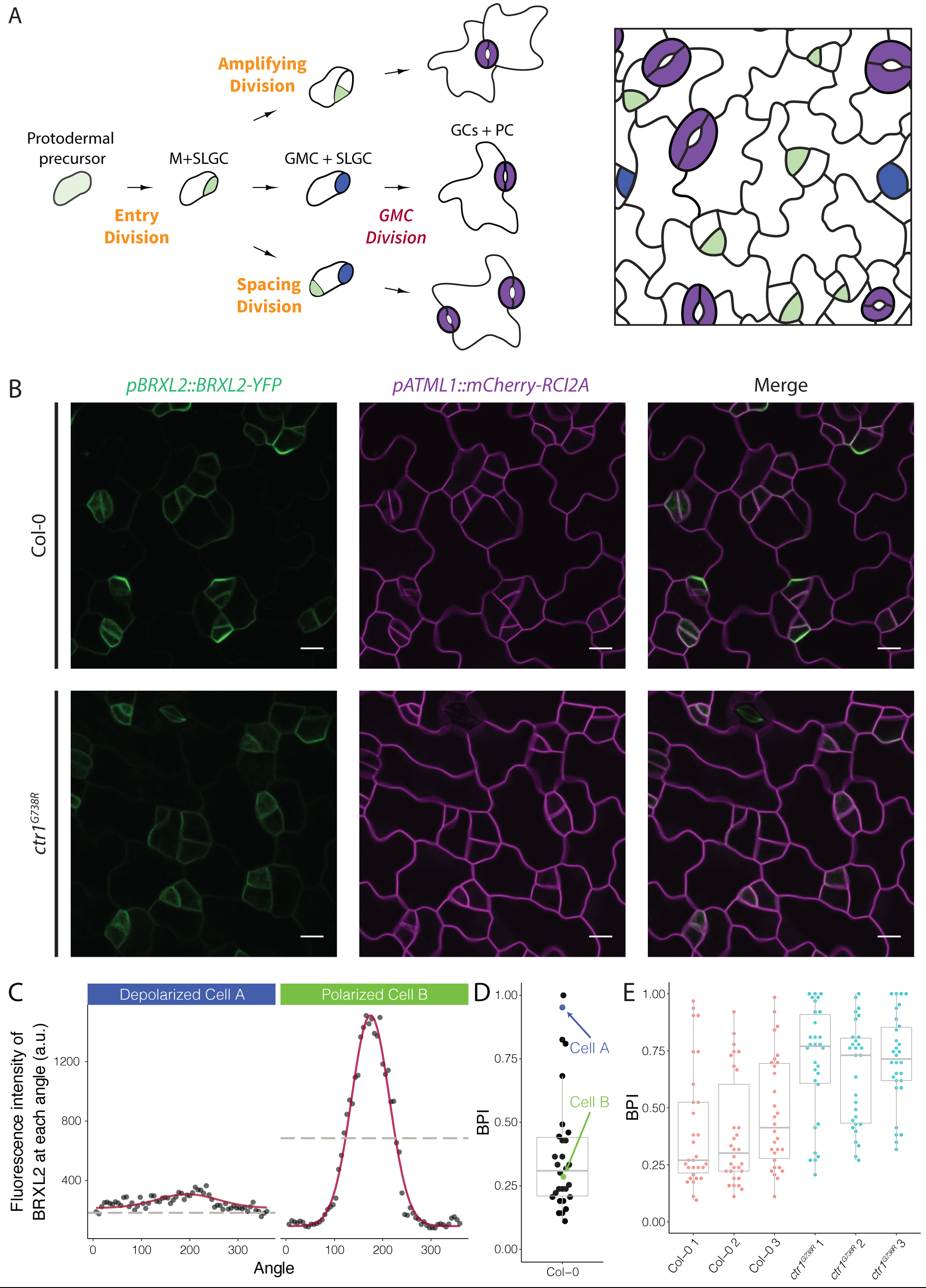
Quantitative analysis of BRXL2-YFP reporter during stomatal lineage divisions reveals reduction in polar localization in the loss of function mutant *ctr1*^*G738R*^. (A) Schematic diagram of stomatal lineage (left) and organization of leaf epidermis (right). From dispersed protodermal precursors, each asymmetric cell division (ACD) produces a small meristemoid (M, green) and a large stomatal lineage ground cell (SLGC, white). The meristemoid can self-renew by undergoing amplifying ACD(s), or differentiate into guard mother cells (GMCs, blue). Each GMC divides symmetrically to produce paired guard cells (GCs, purple). The SLGC can also undergo another ACD (spacing division) or differentiate into a pavement cell (PC). Multiple stomatal lineages are initiated and undergo divisions and differentiation in a dispersed and asynchronized fashion. (B) BRXL2 localization in epidermal cells of 4 dpg Col-0 (top panels) and *ctr1*^*G738R*^ (bottom panels) cotyledons. *pBRXL2::BRXL2-YFP* (left), *pATML1::RCI2A-mCherry* (middle), and merged (right) are shown separately. (C) Output of POME measurement of a cell exhibiting no BRXL2 polarity (cell A, left) and one with polarized BRXL2 (cell B, right). Fluorescence intensity measurements of BRXL2 at each angle are plotted in black dots and the non-linear regression models per each cell are plotted in red. (D) POME quantification of BRXL2 Polarity Index (BPI) in Col-0 (n=30 cells). Each point in graph on right represents a BPI score calculated from the BRXL2 cortical localization pattern of one cell (details in methods and Gong et al. (2020)). (E) Output of polarity measurement (POME) quantification of BRXL2 polarity in 4 dpg Col-0 and *ctr1*^*G738R*^ cotyledons (n=30 cells/genotype, 3 replicates). Scale bar in B, 10 μm.

The following figure supplements are available for figure 1:

**Figure supplement 1.** Molecular description of *ctr1*^*G738R*^ and other alleles, and whole plant phenotypes resulting from *ctr1* mutants, artificial microRNA knockdown, and shoot epidermal-only expression of *CTR1*.

**Figure supplement 2.** Time-lapse imaging of BRXL2 dynamics during stomatal lineage divisions and additional POME quantification of BRXL2 polarity in Col-0 and ctr1G738R.

Several proteins are known to play crucial roles in determining cell fates and division behaviors in the stomatal lineage, including transcription factors (Kanaoka et al., 2008; MacAlister et al., 2007; Ohashi-Ito & Bergmann, 2006; Pillitteri et al., 2007), secreted peptide ligands, and cell surface receptors that mediate cell-cell communication (Bergmann et al., 2004; Hara et al., 2007; Hunt & Gray, 2009; Nadeau & Sack, 2002; Qi et al., 2017; Shpak et al., 2005). The transcription factor SPEECHLESS (SPCH) initiates ACDs, and its expression is maintained briefly in both daughter cells after the ACD (MacAlister et al., 2007). Downstream targets of SPCH include “polarity proteins”: BREAKING OF ASYMMETRY IN THE STOMATAL LINEAGE (BASL), BREVIS RADIX-LIKE 2 (BRXL2) and POLAR (Dong et al., 2009; Lau et al., 2014; Pillitteri et al., 2011; Rowe et al., 2019). These polarity proteins localize to cortical crescents and are required for ensuring the size and fate asymmetries of the ACD (Dong et al., 2009; Houbaert et al., 2018; Pillitteri et al., 2011; Rowe et al., 2019; Zhang et al., 2016; Zhang et al., 2015). Each of these polarity proteins can physically interact with signaling kinases and potentially act as scaffolds to ensure these kinases are active in the appropriate cell types and subcellular locations (Houbaert et al., 2018; Marhava et al., 2018; Zhang et al., 2015). The scaffolded kinases include MITOGEN ACTIVATED PROTEIN KINASES (MAPKs), *Arabidopsis* SHAGGY-LIKE kinases (ATSKs) and AGC kinases—all multifunctional kinases capable of mediating both developmental and environmental signals, thus potentially linking cell polarity to flexible and tunable development.

Despite their potential importance in developmental adaptability, how polarity proteins and their clients initiate and maintain their polarity before and after stomatal ACDs and the mechanistic connection between their polarity and ACDs is largely unknown. From a genetic screen to identify regulators of cell polarity in the stomatal lineage, we found a mutation in *CONSTITUTIVE TRIPLE RESPONSE* (*CTR1*), a core component of the ethylene signaling pathway, that caused overall depolarization of BRXL2. To understand the connection between CTR1 and BRXL2 polarity, and how the change of BRXL2 polarity was affecting leaf development, we created long-term tissue-wide lineage tracing methods and employed a recently developed quantitative polarity analysis tool (Gong et al., 2020) to quantify the polarity of individual cells during cell divisions across cell populations. We discovered that ethylene and glucose signaling, respectively involved in environmental and nutritional pathways, antagonistically regulate the balance of asymmetric and symmetric cell divisions in the stomatal lineage. Additionally, we uncovered a new interaction between the two cells resulting from an ACD, where the temporal dynamics of BRXL2 polarity in an SLGC was linked to the self-renewing capacity of its sister meristemoid. Together, these results reveal previously underappreciated mechanisms that tune stem cell behavior in the stomatal lineage.

## RESULTS

### Loss of CTR1 reduces fraction of stomatal lineage cells with polarized BRXL2-YFP

To identify regulators of cell polarity in *Arabidopsis* stomatal lineage, we performed a microscope-based screen for mutations that affected the subcellular localization of BRXL2. We screened EMS-mutagenized seedlings expressing a native promoter-driven, yellow fluorescent protein (YFP) tagged, BRXL2 reporter (*pBRXL2::BRXL2-YFP*, Figure 1B, top panels). Among recovered mutants, we were particularly intrigued by a line where the BRXL2-YFP signal was mostly depolarized (Figure 1B, bottom panels). Through mapping and cloning by sequencing, we found this recessive mutant contained a G to A mutation in the coding region sequence of *CTR1*. CTR1 is a Raf-like kinase that couples with ethylene receptors and its activity is required to inhibit the downstream ethylene signaling cascade (Huang et al., 2003). The mutation we found is predicted to cause a glycine to arginine substitution at position 738 in the kinase domain (Figure 1—figure supplement 1A) and thus we will refer to this allele as *ctr1*^*G738R*^. Like the previously reported amorphic allele *ctr1-1* (Kieber et al., 1993) and hypomorphic allele *ctr1-btk* (Ikeda et al., 2009; Kieber et al., 1993), *ctr1*^*G738R*^ shows a strong constitutive ethylene response phenotype at the seedling level (Figure 1—figure supplement 1B). The *ctr1*^*G738R*^ mutant also displayed a small leaf phenotype in older plants, similar to *ctr1-1* and stronger than *ctr1-btk*. This phenotype is consistent with leaf epidermal cells undergoing fewer stem-cell like ACDs and an overall decreased number of cells in the leaf epidermis (Figure 1—figure supplement 1C).

To confirm that the disruption of *CTR1* caused the reduction in BRXL2 polarity, we introduced the *pBRXL2::BRXL2-YFP* reporter into the established mutants *ctr1-btk* and *ctr1-1*. We found *ctr1-btk* and *ctr1-1* also caused different degrees of BRXL2 depolarization (Figure 2A-D). We also created a heteroallelic combination of *ctr1*^*G738R*^ with *ctr1-btk*. The F1 progeny of a cross between *ctr1*^*G738R*^ *pBRXL2::BRXL2-YFP* and *ctr1-btk* displayed an intermediate disruption in BRXL2 polarity, as well as an intermediate ethylene response phenotype (Figure 2E). These results confirm *CTR1* as the causal locus, but because *ctr1*^*G738R*^ (like *ctr1-1*) has severe developmental defects, we wanted to eliminate the possibility that the BRXL2 polarity disruption phenotype was a secondary consequence of broad and excessive ethylene signaling. We generated an artificial microRNA (amiRNA) knockdown line targeting *CTR1* under a stomatal lineage-specific promoter (*pTMM::amiRNA-CTR1*). *pTMM::amiRNA-CTR1* lines did not show a strong ethylene response at the seedling stage (Figure 1—figure supplement 1D), but did show reduced polarity of BRXL2 in the stomatal lineage (Figure 2F). Additionally, shoot epidermal-only expression of CTR1 (*pATML1::CTR1*) was sufficient to rescue the reduced BRXL2 polarity phenotype and the small leaf phenotype of *ctr1*^*G738R*^ (Figure 1—figure supplement 1E-H). These rescue results support a direct role of CTR1 in the epidermis, and reinforces previous work showing that the epidermis drives organ growth in leaves.

**Figure 2.**
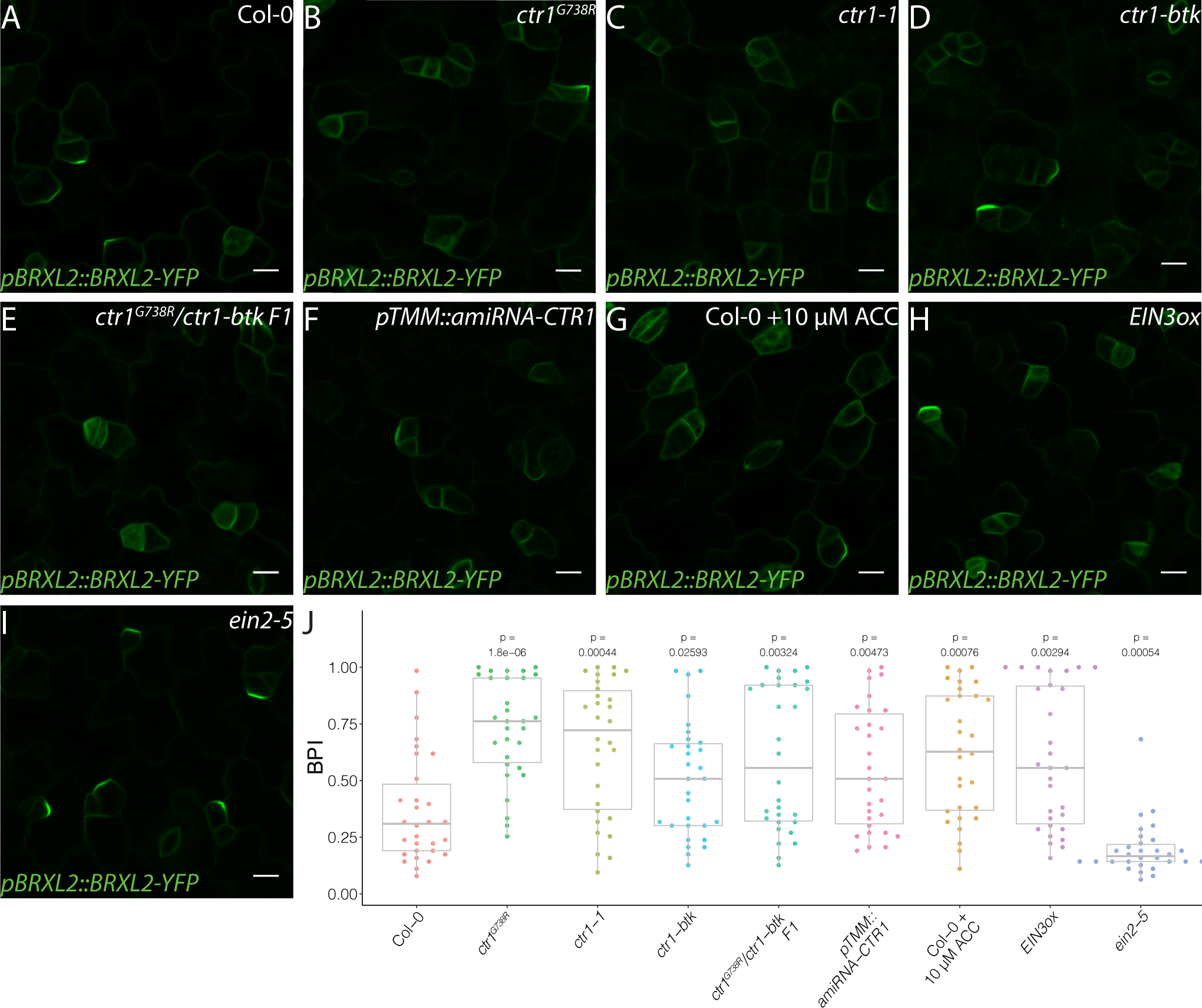
Active ethylene signaling decreases BRXL2 polarity. (A-I) BRXL2 localization pattern in 4 dpg cotyledons from different ethylene mutants or treatment (*pBRXL2::BRXL2-YFP* in green). (A) Col-0, (B) *ctr1*^*G738R*^, (C) *ctr1-1*, (D) *ctr1-btk*, (E) *ctr1*^*G738R*^/*ctr-btk* F1, (F) *pTMM::amiRNA-CTR1*, (G) 10 μM ACC treated Col-0, (H) *EIN3ox*, and (I) *ein2-5*. (J) POME quantifications of BPI from A-H conditions (n=30 cells/genotype). All p-values are calculated by Mann-Whitney test. Scale bars in A-H, 10 μm.

The following figure supplement is available for figure 2:

**Figure supplement 1.** ACC does not affect BRXL2 polarity in ethylene insensitive mutants.

### Time-lapse imaging combined with quantitative polarity analysis reveals nuances of stomatal lineage cell polarity

Among *ctr1* mutant lines, BRXL2-YFP was consistently depolarized in most, but not all, cells. This suggested that CTR1 is not absolutely essential for BRXL2 polarity establishment, and emphasized that to accurately interpret the role of CTR1, we needed a full picture of BRXL2 polarity dynamics throughout the stomatal lineage. We therefore monitored the dynamics of BRXL2-YFP during development of the entire cotyledon over 2 days in 30-minute intervals. Consistent with previous reports (Rowe et al., 2019), a polar crescent of BRXL2 is visible before asymmetric division (Figure 1—figure supplement 2A, top left panels; Supplementary Movie 1), and after division, this crescent is inherited by the large daughter cell (SLGC). Time-lapse, however, revealed two additional features of BRXL2 behavior: first, BRXL2 persists in the SLGC after division for more than 8 hours (Figure 1—figure supplement 2A, top left panels) and second, BRXL2 is still expressed in symmetrically dividing GMCs, but is depolarized in these cells (Figure 1—figure supplement 2A, top right panels). These results suggest a correlation between the degree of BRXL2 polarity and cell identity. Thus, we applied our recently developed <u>po</u>larity <u>me</u>asurement tool (POME, Gong et al., 2020) to quantify the distribution of BRXL2-YFP signal at the periphery of each cell. With POME, the pixel intensity of the BRXL2 reporter and an evenly distributed PM reporter are captured along the entire cell circumference, and their relative distributions were used to compute a “polarity index” (Figure1C, Gong et al., 2020). The <u>B</u>RXL2 <u>p</u>olarity index (BPI) ranges between 0 and 1, where a BPI close to 0 represents a cell with highly polarized BRXL2, and a BPI of 1 represents a cell with completely depolarized BRXL2. As illustrated for Col-0 (WT) in Figure 1D, it is also possible to capture the distribution of BPI measurements in the population of BRXL2-expressing cells. Because of the tight correlation we found between cell identity and BPI, this “snap-shot” population measure can also be used to estimate the ratio of SCDs to ACDs occurring in the leaf epidermis.

We quantified the BPI of Col-0 and *ctr1* cotyledons at 4 days post germination (dpg). As expected, *ctr1*^*G738R*^ cotyledons showed a higher average BPI than Col-0 (Figure 1E), with fewer cells displaying low BPIs. When we specifically analyzed the low BPI cells, we found no significant difference in crescent size or peak height between Col-0 and *ctr1*^*G738R*^ (Figure 1—figure supplement 2B-D). Through additional time-lapse imaging, we demonstrated that, like ACDs in Col-0, stomatal lineage ACDs in *ctr1*^*G738R*^ cotyledons were always preceded by the appearance of polarized BRXL2 in the precursor cell (Figure 1— figure supplement 2A, bottom left panels; Supplementary Movie 2).

These results raised a conundrum: if cells in *ctr1*^*G738R*^ can polarize BRXL2, then how was *ctr1*^*G738R*^ identified and mapped based on a disrupted BRXL2 polarity phenotype? Two potential explanations emerged, each reflecting the dynamic nature of polarity. First, because BRXL2 is polarized in cells undergoing ACDs and depolarized in cells undergoing SCDs, altering the ratios of these division types could decrease the proportion of cells with polarized BRXL2. This change might be detected as a population-level BRXL2 polarity decrease during the screen, and suggests that the role of *CTR1* is primarily to maintain stem cell capacity. Second, we had also observed that BRXL2 polarized crescents did not persist as long after ACDs in *ctr1*^*G738R*^ compared to Col-0 (Figure 6A), and this reduction of BRXL2 persistence would also result in the appearance of relatively fewer polarized cells when observing BRXL2 polarity at a single time point. This result suggests several possible models of CTR1 action in regulating RBXL2 polarity and ACD, but also raises questions about post-divisional functions of polarity factors and how polarity is maintained.

### Disruption of CTR1 results in diminished stem-cell capacity

To dissect the relationship between BRXL2 behavior during ACDs with stem cell capacity and cell fate (stomata and pavement cells) determination, we calculated the stomatal index (SI, ratio of stomata to all epidermal cells) of fully developed cotyledons of Col-0 and *ctr1* mutants at 14 dpg. Compared to Col-0, the SIs of different *ctr1* mutants were significantly elevated (Figure 3A-B, E, and Figure 3—figure supplement 1A-C). Because of the flexible trajectory of the stomatal lineage (Figure 1A), increased SI has several possible origins: (1) an increase in cells entering the stomatal lineage (entry division), (2) an increase in secondary entry via SLGC spacing divisions, or (3) a decrease in meristemoid self-renewal by ACDs (amplifying divisions). To evaluate the contributions of these possibilities, we developed a whole-leaf-based lineage tracing method. We tracked the developmental progression of all epidermal cells in Col-0 and *ctr1*^*G738R*^ cotyledons within a 48hr time window (3 dpg to 5 dpg) and captured the developmental progression of more than 500 pairs of meristemoids and SLGCs per genotype (Figure 3F). Strikingly, the percentage of amplifying ACDs from *ctr1*^*G738R*^ cotyledons was significantly reduced, while the percentage of spacing ACDs was similar to Col-0 (Figure 3G). When individual meristemoids were tracked, we found many underwent fewer rounds of amplifying ACDs during 48 hours in *ctr1*^*G738R*^ compared to Col-0 (Figure 3H).

**Figure 3.**
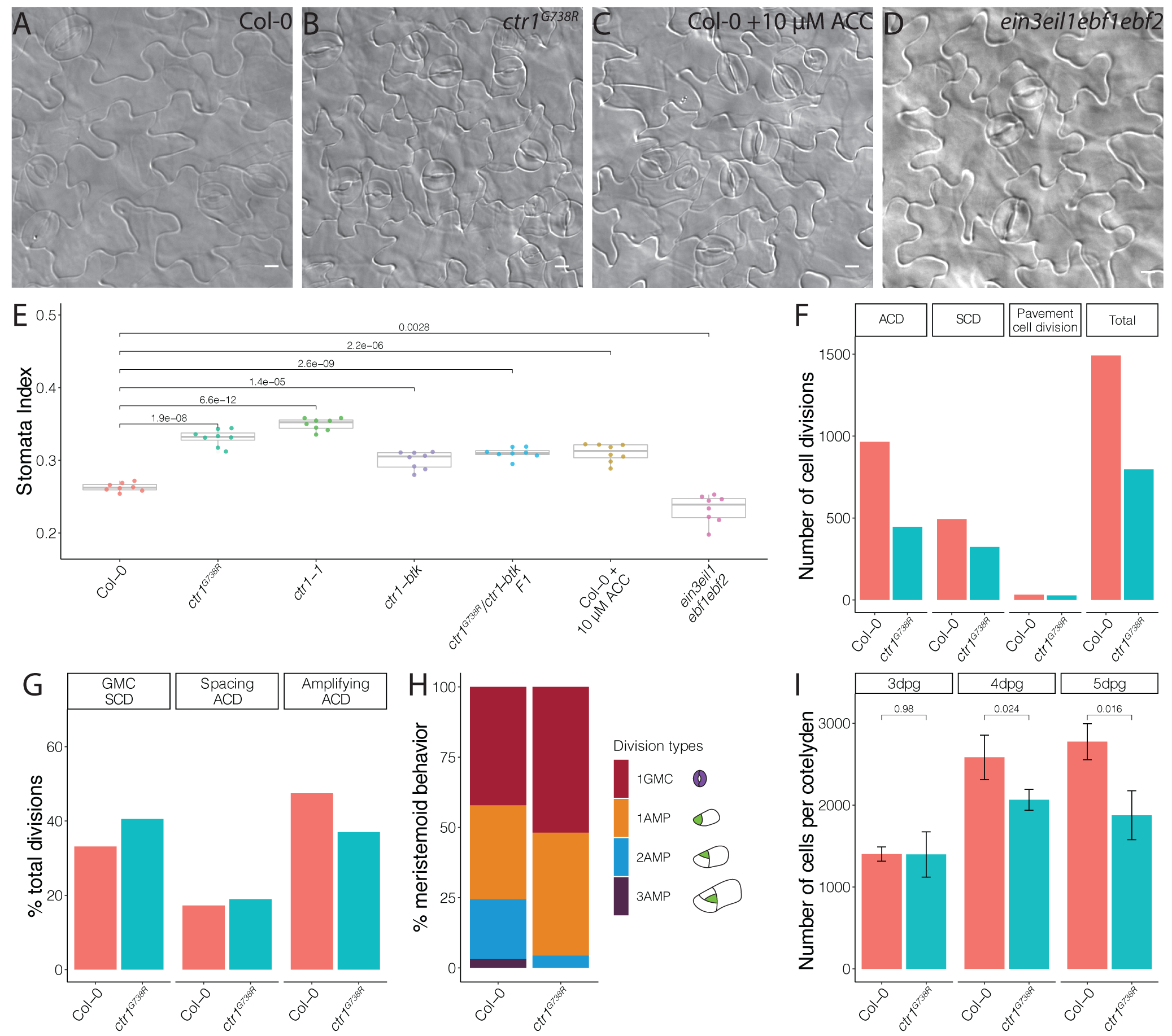
Ethylene signaling modulates the SCD/ACD balance to regulate leaf epidermal development. (A-D) DIC images of (A) Col-0, (B) *ctr1*^*G738R*^, (C) 10 μM ACC treated Col-0, and (D) *ein3eil1ebf1ebf2* 14 dpg cotyledons grown on ½ MS media without sugar. (E) Quantification of the stomatal index of Col-0, ACC treated Col-0, and selected ethylene signaling mutants. (n= 8/genotype). (F-H) Lineage tracing of Col-0 and *ctr1*^*G738R*^ cotyledons. (F) Total numbers of different division types tracked. (G) Percentage of each division type represents among total divisions (H) Cartoon of types of divisions meristemoids undergo, and fraction of each type (n>500 cells/genotype). (I) The epidermal cell number of abaxial cotyledons from 3 dpg to 5 dpg in Col-0 and *ctr1*^*G738R*^. All p-values are calculated by Mann-Whitney test. Scale bars in A-D, 10 μm.

The following figure supplement is available for figure 3:

**Figure supplement 1.** Additional phenotypes of cotyledon epidermis of ethylene signaling mutants.

Fewer amplifying ACDS should result in fewer pavement cells; and this was confirmed by comparing total cell numbers of whole Col-0 and *ctr1*^*G738R*^ cotyledons at 3 dpg, 4 dpg, and 5 dpg. At 3 dpg, *ctr1*^*G738R*^ and Col-0 cotyledons had similar numbers of cells, suggesting a similar level of stomatal entry divisions. However, by 5 dpg, Col-0 had accumulated about 30% more cells than *ctr1*^*G738R*^ (Figure 3I). Together, these results suggest that meristemoids prematurely exit stem cell divisions in *ctr1*^*G738R*^ plants, thereby elevating the SCD/ACD ratio, and ultimately generating fewer epidermal cells, a higher SI, and smaller leaves.

### Ethylene signaling regulates polarity protein complex and stomatal lineage development

CTR1 is best known as a negative regulator of ethylene signaling, but it also is enmeshed in cross-talk with other signaling pathways. To test whether ethylene, in general, affects stomatal lineage ACD/SCD decisions, we treated *pBRXL2::BRXL2-YFP* (Col-0) seedlings with the ethylene precursor 1-aminocyclopropane-1-carboxylic acid (ACC). 10 μM ACC treatment significantly increased average BPI at 4 dpg and SI at 14 dpg relative to mock treated controls (Figure 2G, J, Figure 3C, E). Upon ethylene reception, one of the core elements of the ethylene pathway, ETHYLENE INSENSITIVE 2 (EIN2), is cleaved, translocates to the nucleus, and activates the transcription factors EIN3 and its homolog ETHYLENE INSENSITIVE LIKE 1 (EIL1) mediated ethylene signaling (Alonso et al., 1999; An et al., 2010; Chang et al., 2013; Chao et al., 1997; Qiao et al., 2012; Wen et al., 2012). CTR1 acts as a negative regulator by phosphorylating EIN2 and preventing its cleavage (Qiao et al., 2012; Wen et al., 2012). In plants overexpressing EIN3 (*EIN3ox*) (Chao et al., 1997), the average BPI and SI were higher than the average BPI and SI in WT (Figure 2H, J, Figure 3E, and Figure 3—figure supplement 1D). Conversely, a slight decrease of BPI and SI was observed in the ethylene insensitive mutants *ein2-5* (Alonso et al., 1999) and the quadruple mutant *ein3eil1ebf1ebf2*, where EBF (EIN3-binding F-BOX) proteins were eliminated to approximate complete lack of ethylene response (An et al., 2010, Figure 2I-J, Figure 3D-E). To test whether CTR1 acts primarily through the EIN2/EIN3 core pathway to regulate stomatal stem cell divisions, we treated *ein2-5* single mutants and *ein3eil1ebf1ebf2* quadruple mutants with 10 μM ACC for 4 days. In these ethylene insensitive backgrounds, ACC treatment did not affect BRXL2 polarity (Figure 2—figure supplement 1), suggesting CTR1 acts via the canonical ethylene signaling pathway to modulate meristemoid division behavior, regulate stomatal density, and limit leaf growth.

These genetic and pharmacological perturbations of ethylene signaling indicate that ethylene signaling disrupts BRXL2 polarity primarily through shifting cell identity from meristemoids (which polarize BRXL2) to GMCs (which do not). These results demonstrate how a systemic signal alters tissue level development and suggest that BPI may be an easily scorable proxy for stem-cell potential, a phenotype that was tedious and labor-intensive to measure previously (Vaten et al., 2018).

### Glucose signaling antagonizes ethylene signaling and enhances amplifying divisions

For BPI to be generally useful as an estimate of stomatal stem-cell potential, we needed to identify situations where stem-cell potential was enhanced to provide a counterpoint to the repression we observed in *ctr1*. Previous studies found ethylene and sugar signaling can act in opposition, and so we hypothesized that if sugar signaling also acted in the stomatal lineage, altered availability or perception of sugar would be reflected in the BPI (Haydon et al., 2017; Karve et al., 2012; Yanagisawa et al., 2003; Zhou et al., 1998). Indeed, addition of sucrose to the media resulted in a dose-dependent decrease in average BPI in both *ctr1*^*G738R*^ and Col-0 cotyledons (Figure 4A-D, K, and Figure 4—figure supplement 1).

**Figure 4.**
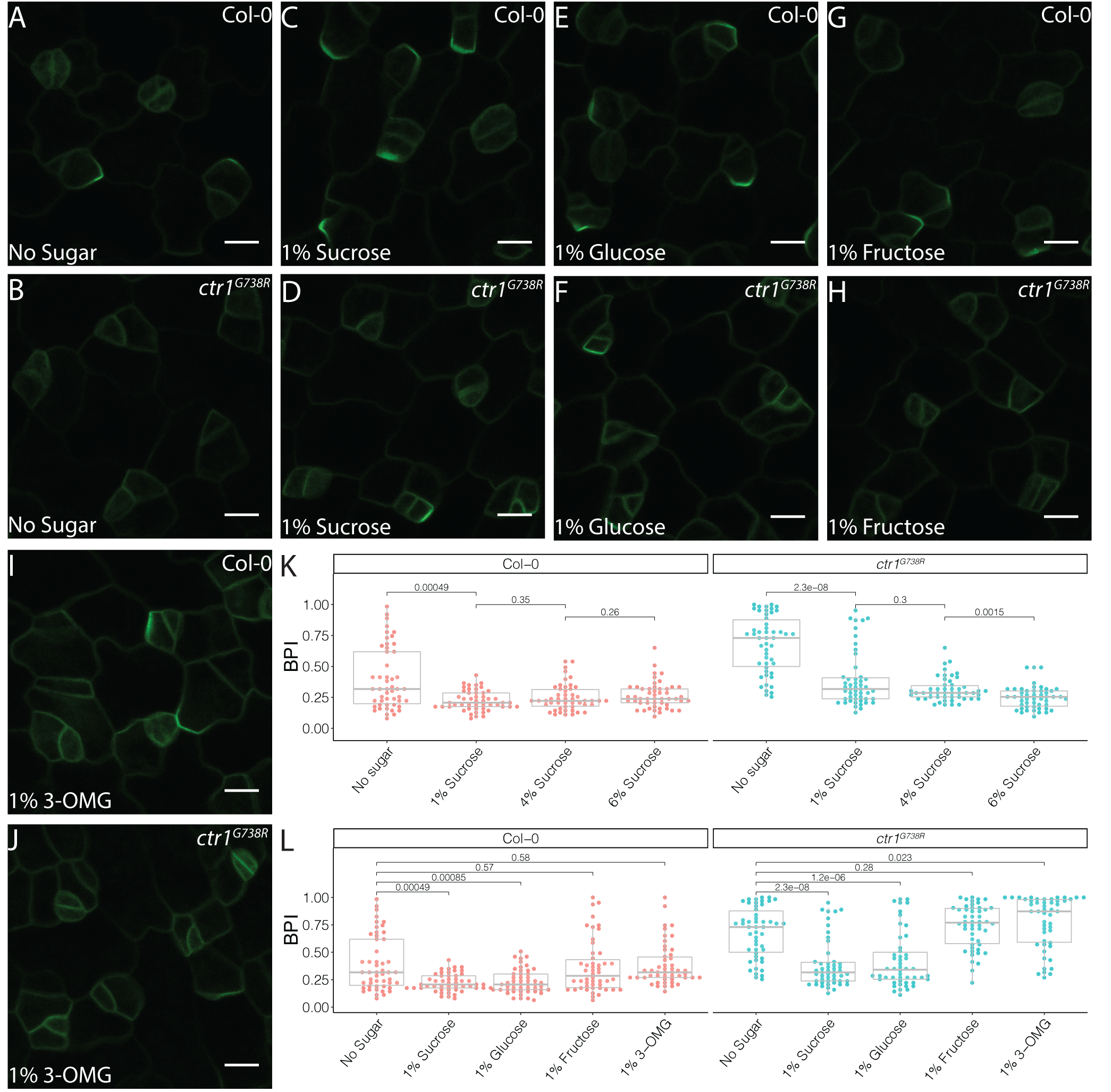
Sucrose and glucose signaling increase BRXL2 polarity. (A-J) BRXL2 localization pattern in 4 dpg cotyledons grown on ½ MS plates with various sugars (*pBRXL2::BRXL2-YFP* in green). (A, C, E, G, and I) Col-0 and (B, D, F, H, and J) *ctr1*^*G738R*^. (A-B) No Sugar, (C-D) 1% sucrose, (E-F) 1% fructose, and (I-J) 1% 3-OMG. (K) BPI quantification of Col-0 and *ctr1*^*G738R*^ grown on ½ MS plates with different sucrose concentrations (n=50 cells/genotype). (L) BPI quantification in 4 dpg Col-0 and *ctr1*^*G738R*^ seedlings growing on ½ MS plates with various sugars (n=50 cells/genotype). The same BPI measurements of Col-0 and *ctr1*^*G738R*^ from no sugar and 1% sucrose treatment are included in K and L for easier visual comparison. All p-values are calculated by Mann-Whitney test. Scale bars in A-J, 10 μm.

The following figure supplement is available for figure 4:

**Figure supplement 1.** BRXL2 localization of Col-0 and *ctr1*^*G738R*^ grown on high levels of sucrose.

The effect of sucrose on BPI could be due to its role as a nutrient or a signal. In *Arabidopsis*, sucrose is the major transport form of sugar and the major energy source for sink tissues where it is further broken down into glucose and fructose to provide energy for cell metabolism, growth, and division. Glucose also acts as a signaling molecule during plant development and responds to environmental cues via a hexokinase (HXK) mediated pathway (Eveland & Jackson, 2012). We therefore tested the ability of metabolizable, non-metabolizable, signaling active, and signaling inactive forms of sugars to change BPI in Col-0 and *ctr1*^*G738R*^. Addition of glucose, but not fructose, decreased average BPI (Figure 4E-H, L), but the glucose signaling inactive analog, 3-O-methyl-D-glucose (3-OMG) (Cortes et al., 2003) failed to alter BPI (Figure 4I-J, L), suggesting that it is glucose signaling, rather than cellular energy status, that affects stomatal lineage progression.

To confirm our model that a decrease in BPI upon glucose treatment reflected an increase in stem cell potential, we tracked division types of all epidermal cells in time courses. Consistent with its ability to decrease average BPI in treated plants, 2% glucose treatment promoted amplifying ACDs in both Col-0 and *ctr1*^*G738R*^ (Figure 5A). Additionally, the SI of *ctr1*^*G738R*^ from 2% glucose growth media also shifted back to the wild-type level (Figure 5B and Figure 5—figure supplement 1), consistent with glucose antagonizing ethylene signaling and boosting the ability of meristemoids to undergo amplifying ACDs.

**Figure 5.**
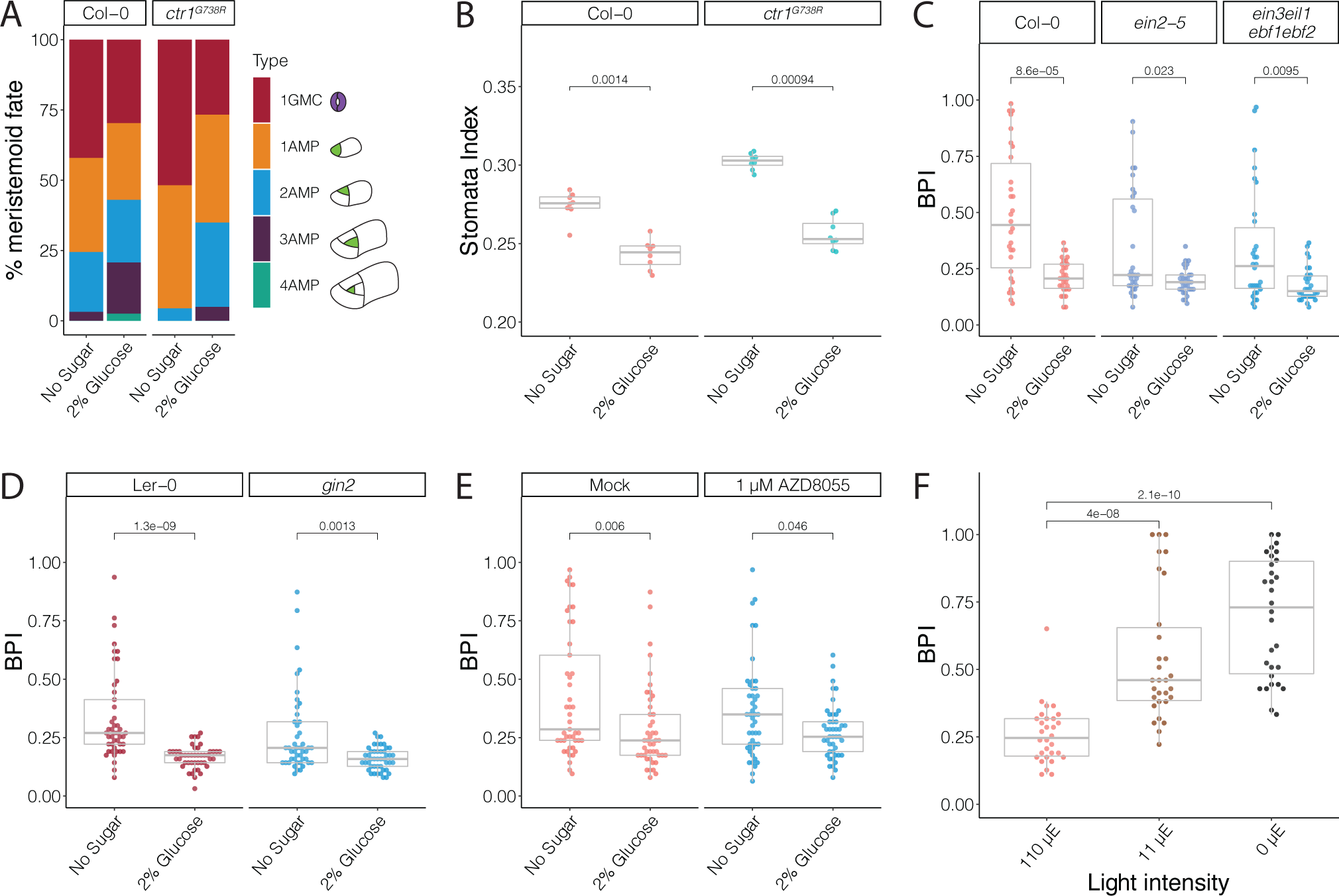
Glucose promotes amplifying ACDs independent of ethylene signaling and HXK1- or TOR-mediated glucose signaling pathways. (A) Cartoon of types of divisions meristemoids undergo, and fraction of each type in Col-0, *ctr1*^*G738R*^, Col-0 with 2% glucose treatment, and *ctr1*^*G738R*^ with 2% glucose treatment (n>500 cells/condition). Data for the no sugar condition of Col-0 and *ctr1*^*G738R*^ are also reported in Figure 3H. (B) Stomatal index of 14 dpg Col-0 and *ctr1*^*G738R*^ growing on ½ MS plates or 2% glucose ½ MS plates (n=8/condition). (C) BPI quantification of 4.5 dpg abaxial cotyledons of Col-0, *ein2-5*, and *ein3eil1ebf1ebf2* grown on ½ MS plates or 2% glucose ½ MS plates (n=30 cells/genotype). (D) BPI quantification of 4 dpg abaxial cotyledons of Ler-0 and *gin2* grown on ½ MS plates or 2% glucose ½ MS plates (n=45 cells/genotype). (E) BPI quantification in 4 dpg Col-0 treated with TOR inhibitor AZD8055 and/or 2% glucose (n=30 cells/genotype). (F) BPI quantification of true leaves in 9 dpg Col-0 seedlings 110 μE normal light condition for 7 days and then transferred to different light intensity conditions for 48 hours (n=30 cells/genotype). All p-values are calculated by Mann-Whitney test.

The following figure supplements are available for figure 5:

**Figure supplement 1.** DIC images of cotyledons from seedlings grown on ½ MS media with or without 2% glucose.

**Figure supplement 2.** BRXL2 localization pattern in Col-0 and ethylene insensitive mutants under different light and sugar treatment regimes.

**Figure supplement 3.** Glucose control of BRXL2 polarity is independent of HXK1 and TOR signaling.

**Figure 6.**
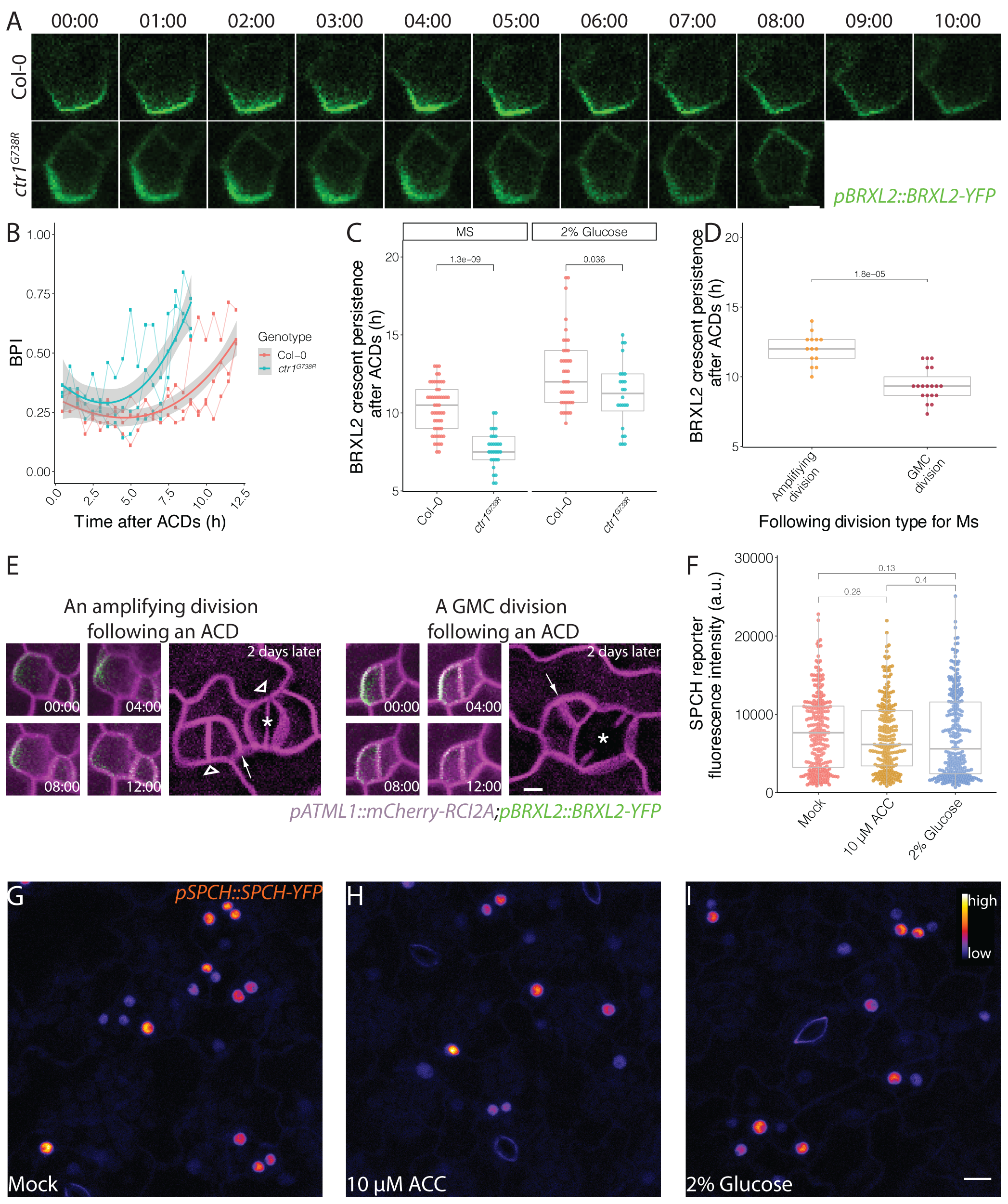
Post-division persistence of BRXL2 polarity is associated with meristemoid fate determinacy. (A) Time-lapse images of Post-ACD BRXL2 dynamics from SLGCs in Col-0 and *ctr1*^*G738R*^ cotyledons (3 dpg). 00:00 (hours: minutes) marks cell plate formation. (B) Quantification of BRXL2 polarity dynamics after ACDs in Col-0 and *ctr1*^*G738R*^. Individual measurement per each cell (n = 3 cells per genotype and 25 timepoints per cell) shown in thin lines and the respective trend per each genotype with 0.95 confidence interval is indicated as the thick line with gray band. (C) Persistence of BRXL2 post-ACD in SLGCs from 3 dpg Col-0 and *ctr1*^*G738R*^ grown in ½ MS media or ½ MS media with 2 percent glucose. (D) Relationship between persistence of BRXL2 in SLGCs and behavior of their meristemoid sisters. (E) Examples of division behaviors quantified in (D). Time-lapse analysis of BRXL2 polarity in 3 dpg Col-0 cotyledons followed by lineage tracing. BRXL2 was imaged every 40 minutes for 16 hours, then plants returned to ½ MS plate for 48 hrs, then re-imaged to capture divisions and fate of the BRXL2-expressing cells. Different division types are marked with asterisks (GMC division), triangles (amplifying division), and arrows (spacing divisions). (F-I) Evidence that ethylene and glucose signaling does not affect SPCH level in individual stomatal lineage cells. F) Quantification of SPCH reporter fluorescence intensity at 4 dpg in (G) mock, (H) 10 μM ACC, and (I) 2% glucose treated Col-0 cotyledons (n = 3 cotyledons/treatment; n>120 cells/treatment). Lookup table Fire is used to false color SPCH reporter intensity (color key in figure). All p-values are calculated by Mann-Whitney test. Scale bars in A and E, 5 μm; I, 10 μm.

The following figure supplement is available for figure 6:

**Figure supplement 1.** Additional characterizations of BRXL2 and SPCH dynamics during ACDs.

### Glucose control of stomatal differentiation is independent of ethylene signaling, HXK1 signaling, or TOR signaling

The relationship between glucose and ethylene signaling has been explored in detail in other contexts, leading to the model that active HKX1-mediated glucose signaling promotes EIN3 degradation and reduces ethylene signaling activity (Yanagisawa et al., 2003). To test if glucose regulates stomatal divisions through EIN3 inhibition, we compared BPIs in the ethylene insensitive mutants *ein2-5* and *ein3eil1ebf1ebf2* grown with and without glucose in the media. Both mutants had significantly reduced BPIs with 2% glucose at 4.5 dpg (Figure 5C and Figure 5—figure supplement 2A-F), suggesting that the influence of glucose in stomatal divisions is independent of the core components of ethylene signaling.

To test another mechanism by which glucose signaling is connected to stomatal lineage development, we quantified the BPI in previously established HXK1 loss of function mutants, *hxk1-3* (Huang et al., 2015) and *gin2* (Moore et al., 2003), in response to glucose treatment. When treated with 2% glucose, both mutants exhibited a decrease in average BPI similar to their corresponding wild-type controls, Col-0 and Ler-0 (Figure 5D and Figure 5—figure supplement 3A-J), suggesting that glucose’s regulation in stomatal development is not mediated by HXK1. The Target of Rapamycin signaling pathway (TOR) has been suggested to act downstream of sugar signaling as a major controller of plant growth-related processes, including meristem proliferation, leaf initiation, and cotyledon growth (Li et al., 2017; Rexin et al., 2015; Xiong et al., 2013). Because TOR null mutants are lethal and land plants are largely insensitive to rapamycin, we treated *pBRXL2::BRXL2-YFP* expressing seedlings grown in media with and without 2% glucose with 1 μM AZD-8055, an ATP-competitive inhibitor of TOR (Montane & Menand, 2013), for 2 days. Despite the presence of AZD-8055, glucose still decreased the average BPI (Figure 6E and Figure 5—figure supplement 3K-N). Therefore, despite clear evidence that glucose signaling can influence stomatal lineage behaviors, we have been unable to link this regulation to known HXK1 or TOR mediated sugar signaling pathways.

### Stomatal BPI can respond to physiological depletion of sugars

By experimentally adding sugars we could modulate BPI and stomatal divisions. The critical question then becomes whether this is biologically relevant—do *Arabidopsis* leaf epidermal cells sense endogenous levels of glucose or sucrose and adjust the stomatal lineage to create leaves of appropriate cellular composition? To test this, we adapted an approach used in (Moraes et al., 2019), and reduced light intensity to limit the photosynthetic rate in seedlings. Because sugar is the primary product of photosynthesis, this experimental procedure serves to exhaust endogenous sugars in leaves. We transferred 7 dpg Col-0 seedings from our regular high-light intensity (110 μE) growth condition to low light conditions (10 μE or 0 μE). After 48 hours, seedlings grown in low light conditions showed increased BPIs in their leaves (Figure 5F and Figure 5—figure supplement 5G-I), indicating fewer ACDs took place. Thus, the decrease of ACDs seen in low light-grown seedlings is consistent with sucrose or glucose concentration in the leaf providing feedback to coordinate leaf development with photosynthesis.

### Post-division BRXL2 crescent is associated with meristemoid fate determinacy

In previous sections, we showed that CTR1, ethylene, and sugar signaling regulate the balance between stomatal lineage cells undergoing proliferative ACDs and differentiating SCDs, where the connection to BRXL2 polarity is largely indirect. However, in our examination of the *ctr1*^*G738R*^ mutant, we also noticed that polarized BRXL2 was in a smaller proportion of SLGCs than in Col-0, and suspected that BRXL2 was less persistent in these cells. We investigated this observation more rigorously by measuring the persistence of the polarized BRXL2 crescent in multiple genotypes under different growth conditions. From these time-lapses, we noticed that the BRXL2 crescent was significantly less persistent in *ctr1*^*G738R*^ than in Col-0 (Figure 6A-B and Figure 6—figure supplement 1A) and glucose increased persistence of BRXL2 crescent in both *ctr1*^*G738R*^ and Col-0 seedlings (Figure 6C). Together, the observations that meristemoids in *ctr1*^*G738R*^ underwent fewer amplifying ACDs (Figure 3H) and glucose promoted amplifying ACDs in both Col-0 and *ctr1*^*G738R*^ (Figure 5A) indicates a positive correlation between the persistence of the post-division BRXL2 polarity complex in the SLGC and the self-renewal capacity of the meristemoid (the SLGC sister derived from the previous ACD).

The potential non-cell-autonomous effect of BRXL2 persistence suggests that either there is communication between a SLGC and its meristemoid sister to influence the meristemoid’s cell fate, or, that by monitoring BRXL2, we witnessed a differentiation event already specified in their mother cell; for example, a change in SPCH activity that distinguishes cells with higher self-renewal potential and cells that will undergo their final ACDs.

SPCH initiates all the ACDs in the stomata lineage (MacAlister et al., 2007), and SPCH directly binds the promoters of *BRXL2* and *BASL* in ChIP-Seq studies (Lau et al., 2014). BRXL2 crescent persistence, and the change of that persistence in response to ethylene and glucose treatment, therefore, might be directed by quantitative changes in SPCH expression in the cells undergoing ACD. We quantified SPCH protein levels in individual cells from 10 μM ACC, 2% glucose, and mock treated Col-0 cotyledons by measuring the fluorescent intensity of a functional SPCH reporter (*pSPCH::SPCH-YFP* in *spch3*, Vaten et al., 2018). We found no significant change in SPCH-YFP levels upon treatment with either ACC or glucose (Figure 6F-I). We also examined SPCH reporter dynamics during ACDs in these different treatment conditions, and found no obvious changes in SPCH reporter peak intensity, onset of expression, or post-divisional persistence (Figure 6— figure supplement 1C).

If BRXL2 crescent persistence in an SLGC is not coupled to the upstream (SPCH) transcriptional response, then this persistence is unlikely to be reflecting a decision made pre-ACD in the mother cell. To correlate BRXL2 crescent persistence post-ACD with the subsequent divisions and fates of the daughters, we performed detailed time-lapse imaging (16hr, 40 min intervals) followed by a 48hr time-course in Col-0. We found that the post-ACD BRXL2 crescent persistence in SLGCs was predictive of sister meristemoid behavior: BRXL2 crescents persisted, on average, two hours longer when meristemoids underwent amplifying ACDs than when meristemoids divided symmetrically (Figure 6D-E). Interestingly, BRXL2 was not predictive of the behavior of the cell in which it resides: there was no significant difference between persistence in SLGCs that underwent spacing ACDs and those that did not (Supplementary Figure 1B). Together, these results suggest there must be communication between the sister cells resulting from an ACD. Whether this communication is through secreted ligand/cell-surface receptor interactions, through other chemical signals, or even through a mechanical signal induced by differential growth is beyond the scope of this report, but these and other plausible signaling scenarios are summarized in Figure 7.

**Figure 7.**
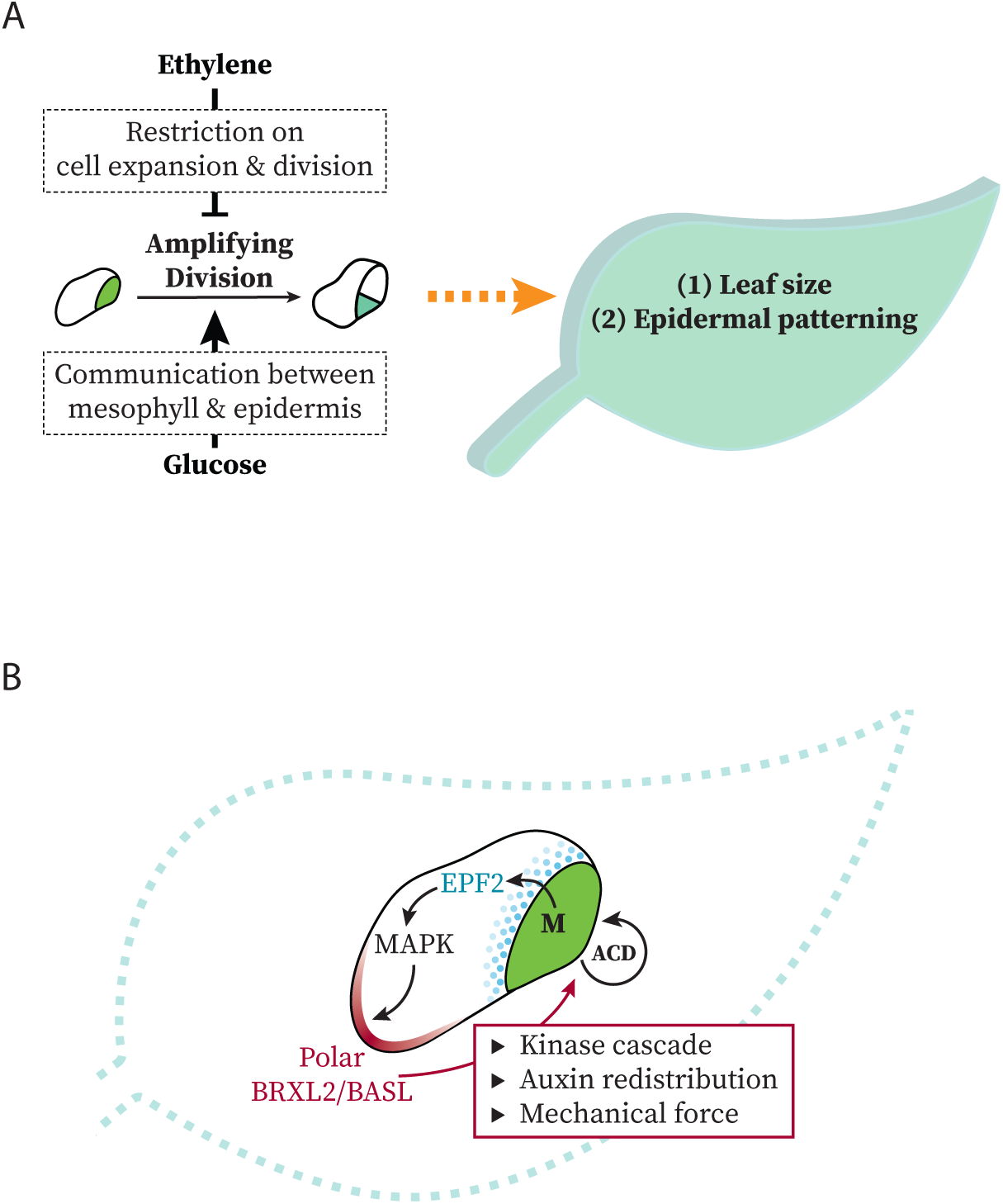
Models at organ and cell scales for connection between systemic signaling, cell polarity, stomatal stem-cell potential and leaf growth. (A) Schematic representation of the regulation of ethylene and glucose on meristemoids’ amplifying divisions. Potential regulatory mechanisms used by these signals are illustrated in dashed box. (B) Schematic representation of modes of communication possible between SLGCs and meristemoids during polarity and cell division control.

## DISCUSSION

Plants respond to environmental stimuli by modifying their development. As the major organs of photosynthesis, leaves must regulate their size, position and gas-exchange capacity to adapt and compete. Using a combination of genetics, live-imaging, quantitative image analysis and lineage tracing of the simple, yet flexible, *Arabidopsis* stomatal lineage, we have been able to link ethylene and sugar signaling to the self-renewing capacity of epidermal stem cells. The immediate readout of ethylene and glucose antagonism is a shift in population of cells expressing polarized vs. depolarized BRXL2, and the ultimate readout is a change in the size and the cell type composition of the leaf (Figure 7A). In addition, by developing methods to systematically monitor fluorescent marker dynamics (POME, Gong et al., 2020) and to then track cell divisions and differentiation in the same cells, we uncovered a surprising correlation between the ACD potential of meristemoids and the persistence of polarized BRXL2 in SLGCs (Figure 7B), suggesting active communication and coordination between these sister cells.

Finding *CTR1* alleles in screens for ACD regulators was unexpected, and we considered whether there was a true role for ethylene signaling, or whether we were seeing the effect of altering *CTR1* on other pathways. Genetic and pharmacological experiments (Fig 2 and 3), however, confirmed the participation of multiple ethylene signaling components in regulating stomatal lineage ACDs. Ethylene is considered an “aging” and a “stress” hormone (Iqbal et al., 2017; Schaller, 2012). Additionally, ethylene also regulates many different aspects of plant development, often through its crosstalk with auxin (Muday et al., 2012; Strader et al., 2010; Vaseva et al., 2018; Wen et al., 2012). In both root and leaf epidermal cells, for example, ethylene promotes local auxin biosynthesis. Elevated auxin level then inhibits cell expansion (Vaseva et al., 2018) and in turn induces expression of ethylene biosynthesis genes (Abel et al., 1995; Tsuchisaka & Theologis, 2004). Is ethylene’s regulation on stomatal lineage cell divisions also through its feedback loop with auxin? In the stomatal lineage, auxin has also been shown to influence cell division and differentiation in the stomatal lineage, but there are some conflicting results, with Le et al. (2014) suggesting that auxin promotes meristemoid ACDs and Balcerowicz et al. (2014) concluding that auxin inhibits ACDs. The interplay of auxin and ethylene and the influence of auxin on all types of stomatal lineage divisions and fate transitions is beyond the scope of this manuscript, but will be an exciting future direction and may be answered more definitively by new tools like engineered TIR1 (Uchida et al., 2018) expressed in specific stomatal lineage cell types.

We did pursue the cross-talk between ethylene and sugar signaling, and found that higher levels of each resulted in opposite effects on BRXL2 persistence and on stomatal lineage ACDs. Introduction of 2% glucose to growth media increased the stem-cell like ACDs even in ethylene insensitive mutants indicating that the epidermis is likely perceiving these signals independently. An effect of sugars on stomatal production was recently reported (Han et al., 2020) and while their experimental conditions differ substantially from ours, both studies concur that sugars promote stomatal development. What information is sugar providing? Glucose and sucrose are the major products of photosynthesis produced in mesophyll cells, and recent work demonstrates multiple modes of communication between mesophyll and epidermis to coordinate these tissues for optimal growth and gas exchange (Baillie & Fleming, 2020; Dow et al., 2017; Sugano et al., 2010). It is attractive to consider mesophyll-derived sugar signaling as a way to promote growth and stomatal production, and future experiments could be designed to trace the source of sugar perceived by the stomatal lineage.

Perhaps our most interesting and surprising result was that temporal variations in BRXL2 persistence correlated with ACD potential. In particular, although BRXL2 is expressed in SLGCs, it was the behavior of the sister meristemoid that was affected. This made us consider what properties of the meristemoid are most important for that cell’s behaviors (Figure 7B). Previous work considered the expression of the transcription factor SPCH as the key to modulating stomatal ACDs (Lau et al., 2018; Simmons et al., 2019; Vaten et al., 2018; Zhang et al., 2016). However, while SPCH is necessary for divisions, we did not find that glucose or ethylene signaling had a dose-dependent effect on SPCH levels. On the other hand, meristemoids vary in cell size. Additionally, ethylene inhibits and sugar promotes cell expansion, so one possibility, parallel to the situation in the *C. elegans* germline, is that smaller meristemoids undergo fewer rounds of ACDs before committing to terminal differentiation. This hypothesis also could explain the difference in ethylene’s effect on ACD potential in meristemoids and SLGCs. Since SLGCs are larger than meristemoids, if there is a minimal absolute cell size for the ACD, SLGCs would be most likely above the size threshold, rendering them less sensitive to ethylene’s inhibition of cell expansion.

By what mechanisms could SLGCs influence their sister meristemoid, and how might persistent BRXL2 polarity drive this regulation? Signals from the meristemoid to the SLGC rely on EPIDERMAL PATTERNING FACTOR2 (EPF2) mediated cell-cell signaling that results in higher levels of MAPK signaling in SLGCs (Lee et al., 2015). Elevated MAPK signaling not only inhibits divisions (Bergmann et al., 2004; Lampard et al., 2009), but also enhances BASL polar localization (Zhang et al., 2015). In this scenario, BRXL2 polarity persistence in the SLGC could be a readout of the ACD potential of the sibling meristemoid. We propose, however, the longer maintenance of the polarity domain in the SLGC can also act as the source of a signal because of the relationship between “polarity proteins” BASL, BRXf, and POLAR and the kinases (MAPKs, PAX, BIN2) these proteins scaffold (Houbaert et al., 2018; Marhava et al., 2018; Zhang et al., 2015). For example, plant MAPKs activate numerous target genes, and some could initiate production of mobile signals that would accumulate to biologically active levels over time. BRXf and POLAR have been linked to auxin and brassinosteroid hormone signaling, respectively, and these signals have numerous connections to stomatal lineage progression (Houbaert et al., 2018; Kim et al., 2012; Le et al., 2014). At the same time, BASL and BRXL2 polar crescents mark tissue level polarity fields, and appear to precede local cell outgrowth (Bringmann & Bergmann, 2017; Mansfield et al., 2018). Local cell expansion in SLGCs itself could also serve as a division-promoting signal to the meristemoid, as has been seen in other contexts where such a mechanism is used to maintain tissue integrity among mechanically coupled cells (Hamant & Haswell, 2017). Distinguishing among these relationships will require new tools to specifically alter polarity crescent persistence in an otherwise unperturbed background, but advances in inducible degradation systems (Faden et al., 2016; Sallee et al., 2018) may enable these experiments in the future.

## MATERIALS AND METHODS

**Key resources table**

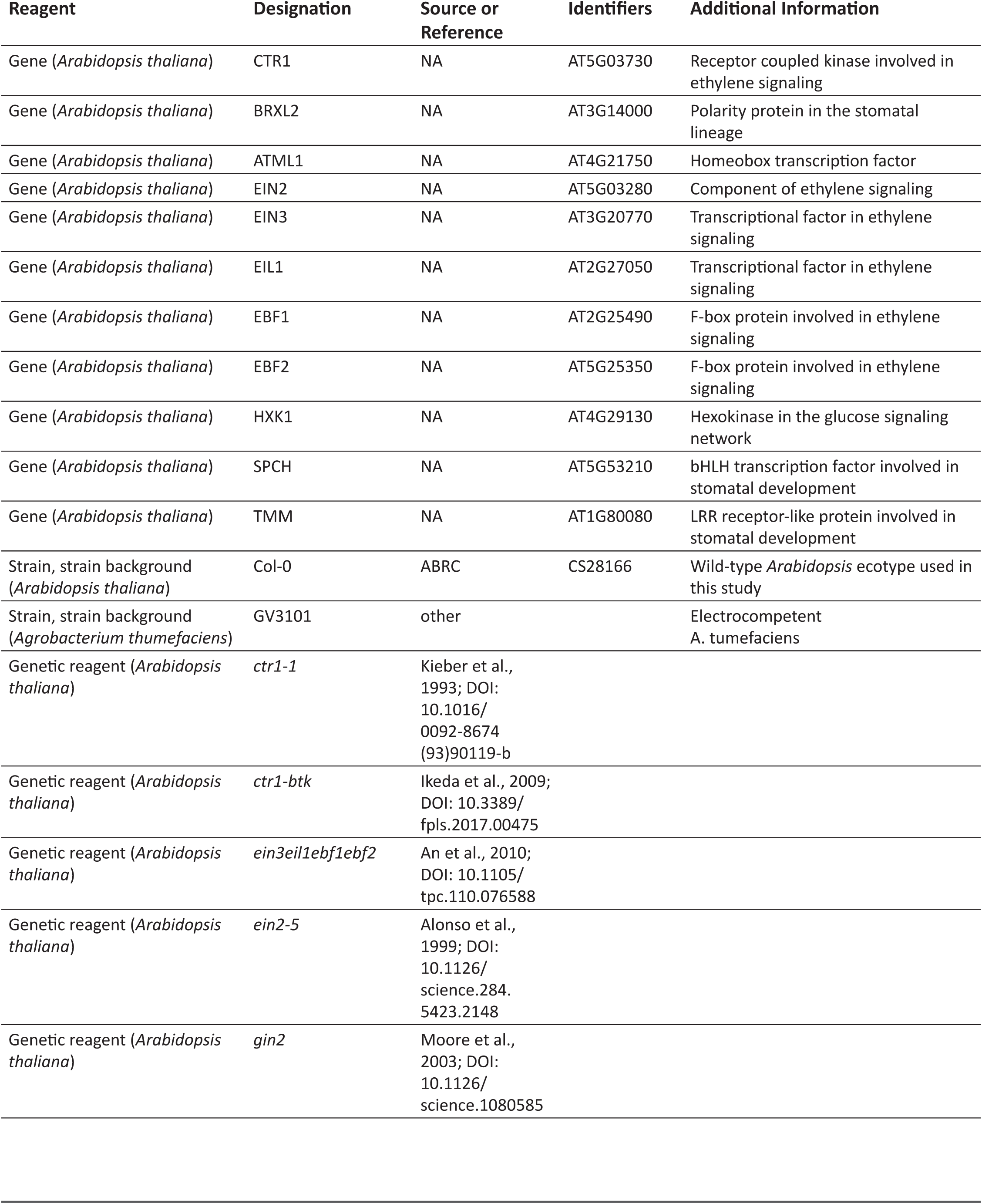
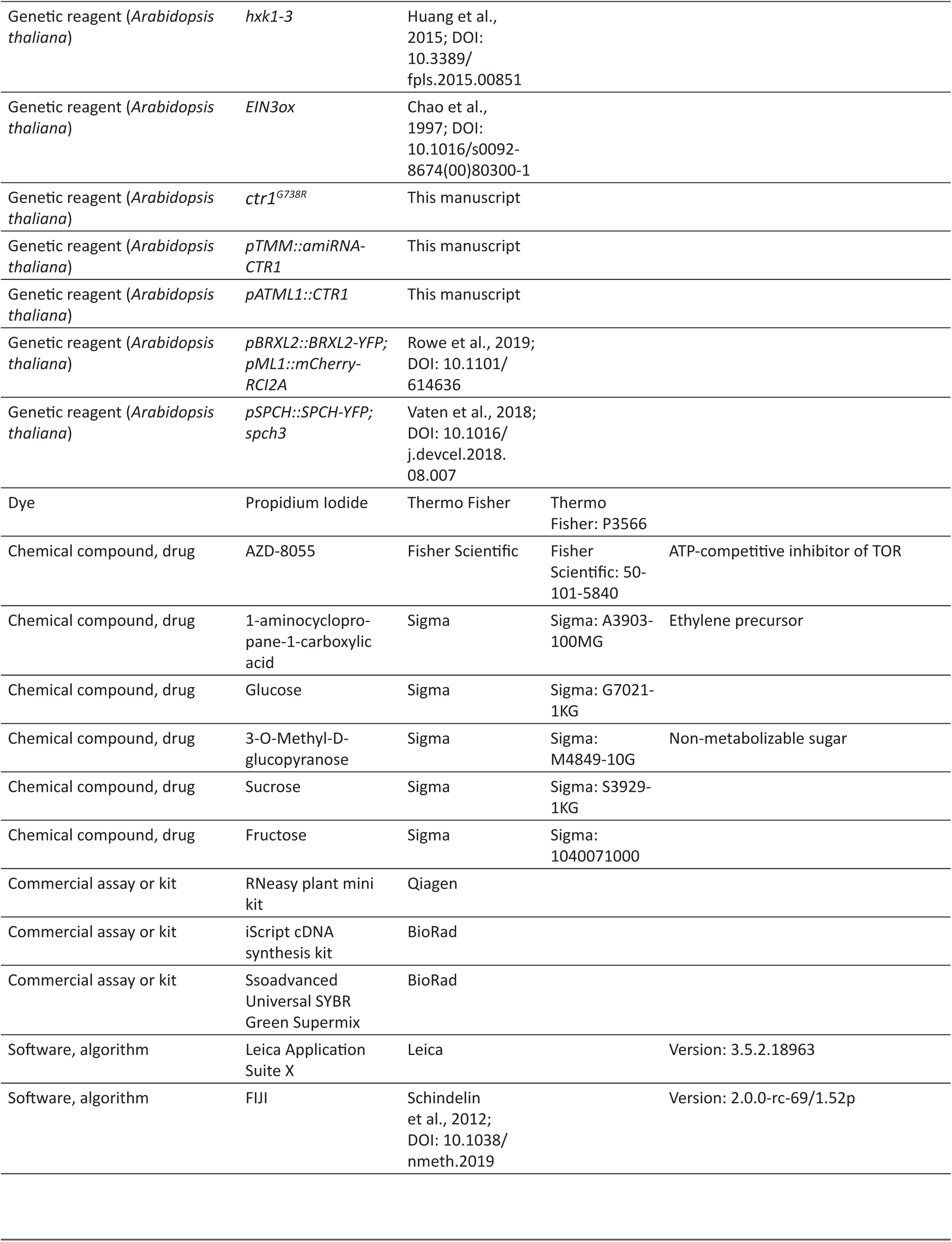
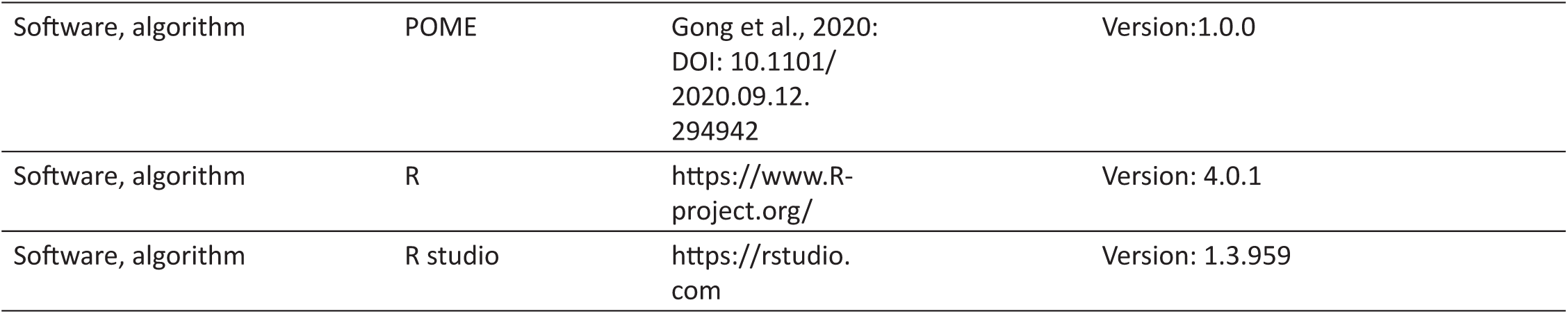

### Plant material and growth conditions

All *Arabidopsis* lines used in this study are in Col-0 background, and sources of previously reported mutants and transgenic lines are listed in the table above. Newly generated lines include *pTMM::CTR1amiRNA* and *pATML1::CTR1*.

All *Arabidopsis* seeds were surface sterilized by bleach or 75% ethanol and stratified for 2 days. After stratification, seedlings were vertically grown on ½ Murashige and Skoog (MS) media with 1% agar for 3-14 days under long-day condition (16h light/8h dark at 22°C) and normal light condition (110 μE) unless noted otherwise.

### Identification and map-based cloning of *ctr1G738R*

Col-0 seeds homozygous for the reporter *pBRXL2::BRXL-YFP* were mutagenized with Ethylmethanesulfonate (EMS). Seedlings from M2 families were screened individually at 3-5 dpg for loss of polar YFP localization on a Leica SP5 confocal microscope. Candidate mutants were then outcrossed to Col-0 and F2 progeny with and without the mutant phenotype (>50 plants of each type) were pooled, sequenced and analyzed as in Wachsman et al. (2017).

### Vector construction and plant transformation

To generate *pTMM::amiRNA-CTR1*, an artificial microRNA sequence targeting *CTR1* was designed with the Web MicroRNA Designer platform (http://wmd3.weigelworld.org) (Schwab et al., 2006), engineered with the pRS300 plasmid with the TMM promoter, and cloned into the binary vector R4pGWB401 (Nakagawa et al., 2008). *pATML1::CTR1* was generated by cloning CTR1 coding sequence from cDNA into pENTR and combining with ML1 promoter sequences in binary vector R4pGWB401. Primers used to generate these two constructs are listed in the primers section below. Transgenic plants were then generated by Agrobacterium-mediated transformation (Clough & Bent, 1998), and transgenic seedlings were selected on ½ MS plates with 50 μM Kanamycin.

### Drug treatments

To assay the influence of different types and concentrations of sugar on BPI or SI, filter sterilized 40% sucrose, 20% glucose, 20% fructose, or 20% 3-OMG water solutions were prepared as stock solutions and added to the sterilized ½ MS media with 1% agar to make ½ MS sugar treatment plates. Similarly, ACC was dissolved in water and filter sterilized to create the 100 mM ACC stock solution. This ACC stock solution was then added to the sterilized ½ MS media with 1% agar to make ½ MS ACC treatment plates. Surface sterilized and stratified *Arabidopsis* seeds were then plated on these plates and vertically grown for 3-14 days under long-day condition for the respective experiments. For AZD-8055 experiments, AZD-8055 was dissolved in DMSO to create 1 mM stock solution. AZD-8055 stock solution or DMSO was then added to the sterilized ½ MS media with 1% agar to make ½ MS AZD-8055 or mock treatment plates. Surface sterilized and stratified *Arabidopsis* seeds were then plated on regular ½ MS plates and vertically grown for 2 days prior to being transferred to AZD-8055 or mock treatment plates and vertically grown 2 more days before image acquisition. For time-lapse experiments, ½ MS solution was supplemented with 2% glucose prior to seedling loading and all flow-through solution contained the same glucose concertation.

### Microscopy and image acquisition

All fluorescence imaging experiments were performed on a Leica SP5 confocal microscope with HyD detectors using 40x NA1.1 water objective with image size 1024*1024 and digital zoom from 1x to 2x.

For time-lapse experiments, 3 dpg seedlings were mounted in a custom imaging chamber filled with ½ MS solution (Davies & Bergmann, 2014). Laser settings for each reporter, except the membrane marker (*pATML1::mCherry-RCI2A*), were adjusted to avoid over-saturation. For the time-lapse experiments reported in this study, there was a 30-45 min interval between each image stack capture. For the reoccurring time-lapse experiments, seedlings were imaged in the time-lapse chamber for 16 hours with the protocol stated above. After imaging, seedlings were removed from the imaging chamber and returned to MS-agar plates (with appropriate supplements) for 8 hours under standard light and temperatures. The same epidermal surface from the same plant was re-imaged using the same time-lapse protocol each successive day from 3-5 dpg. The three sets of time-lapse images were combined together, with the time between recordings noted, to create a time-lapse covering about 64 hours of development.

For time-course (lineage tracing) experiments, where images of the same whole epidermis were acquired every 24 hours from 3 dpg to 5 dpg, each seedling was carefully mounted on a slide with vacuum grease outlining the border of the cover slide. In this setting, vacuum grease provides mechanical support to avoid crushing the seedlings. After each image acquisition, seedlings were carefully unmounted from the slide and returned to the ½ MS plate and normal growth condition until the next image acquisition. For the analysis of these time-course (lineage tracing) experiments, refer to the “Lineage tracing analysis” section below.

All raw fluorescence image Z-stacks were projected with Sum Slices in FIJI unless noted otherwise. For all time-lapse images, drift was corrected using the Correct 3D Drift plugin (Parslow et al., 2014) prior to any further analysis.

For SI counting, seedlings were collected at 14 dpg. Samples were cleared with 7:1 solution (7:1 ethanol:acetic acid), treated with 1 N potassisum hydroxide solution, rinsed in water, and then mounted in Hoyer’s solution. Individual leaves were then imaged with a Leica DM6B microscope with 20x NA0.7 air objective in differential contrast interference mode.

### Image quantification

For POME measurements of BRXL2 polarity in different mutants and treatments, florescence images of BRXL2 and membrane marker or staining in these conditions were acquired with 40x water objective and 2x digital zoom. 3 to 5 individual images were captured from the same region of cotyledon from different seedlings. For each individual image, relative brightness of BRXL2 reporter was used to select the 10 brightest cells, and POME was used to measure BRXL2 distribution along the cell membrane in each of the selected cells. This selection was made as a way to capture cells that had recently divided either symmetrically or asymmetrically, and was the fairest comparison of BRXL2 polarity across different mutants and treatments. With POME, the cortex of each cell was divided in 63 portions and the fluorescence intensity of BRXL2 reporter in each portion was measured and reported. Details of BRXL2 measurement with POME are available in Gong et al. (2020). BPI is defined as the fraction of measurements above the half maximum. To calculate the BPI for each cell, the maximum BRXL2 fluorescence intensity of all 63 measured portions was determined, and the fraction of all measurements with BRXL2 fluorescence intensity above half of this maximum was calculated as the BPI. BPIs of all the measured cells in the same conditions were then grouped, plotted, and compared with other conditions.

For the quantification of BRXL2 polarity persistence, the duration of BRXL2 polarity persistence was counted manually in different genotypes and treatments. For each cell, the beginning of post-ACD BRXL2 polarity was set as the image in which formation of the cell plate was first visible, while the end of post-ACD BRXL2 was set as the first time-frame where BRXL2 polar crescent was no longer detectable. Post-ACD BRXL2 polarity dynamics from three individual cells of Col-0 and *ctr1*^*G738R*^ under no sugar condition were measured with POME (Gong et al., 2020), and the dynamic of the BPI and the normalized crescent amplitude (peak amplitude divided by that at timepoint 0) were quantified and plotted in Figure 6B and Figure 6 —figure supplement 1A.

### Lineage tracing analysis

In each lineage tracing experiment, whole leaf images of the same leaves from different days were grouped together. Each individual cell from the 3 dpg (or any starting time point) was then annotated for divisions and lineage relationship each day after the starting point. BRXL2 reporter presence and polarity level in each cell and in different days was also recorded to help determine the division type, as GMC divisions display depolarized BRXL2 while amplifying and spacing ACDs harbor a polarized BRXL2 crescent. After all the cells on the leaf epidermis were annotated, division behaviors of all cells are summarized in a spreadsheet, from which the total number and the fraction of each division type, fraction of each meristemoid behavior, total number of cells at the starting and ending time points, and any other developmental behaviors of interest were calculated.

### RNA extraction and qRT-PCR analysis

Whole seedlings of Col-0 and *hxk1-3* at 7 dpg were collected. Seven seedlings were grouped into a biological replicate and two biological replicates per genotype were assayed. RNA of each biological replicate was then extracted with RNeasy plant mini kit (QIAGEN) with on-column DNAse digestion. 1ug of total RNA was used to for cDNA synthesis using the iScript cDNA synthesis kit (Bio-Rad). The qPCR reactions were performed on a CFX96 Real-Time PCR detection system (Bio-Rad) with the Ssoadvanced Universal SYBR Green Supermix (Bio-Rad). Three technical replicates were performed per biological replicate. Expression level of *HXK1* was then calculated and normalized to the expression level of the reference gene *UBC21* using the ΔΔCT method.

### Statistical analysis

All statistical analyses in this manuscript were performed in RStudio. Unpaired Mann-Whitney tests were conducted to compare two data samples with compare_means function from the ggpubr package (Kassambara, 2020). For all graphs, p-values from the unpaired Mann-Whitney tests were directly labelled on these graphs.

### Primers

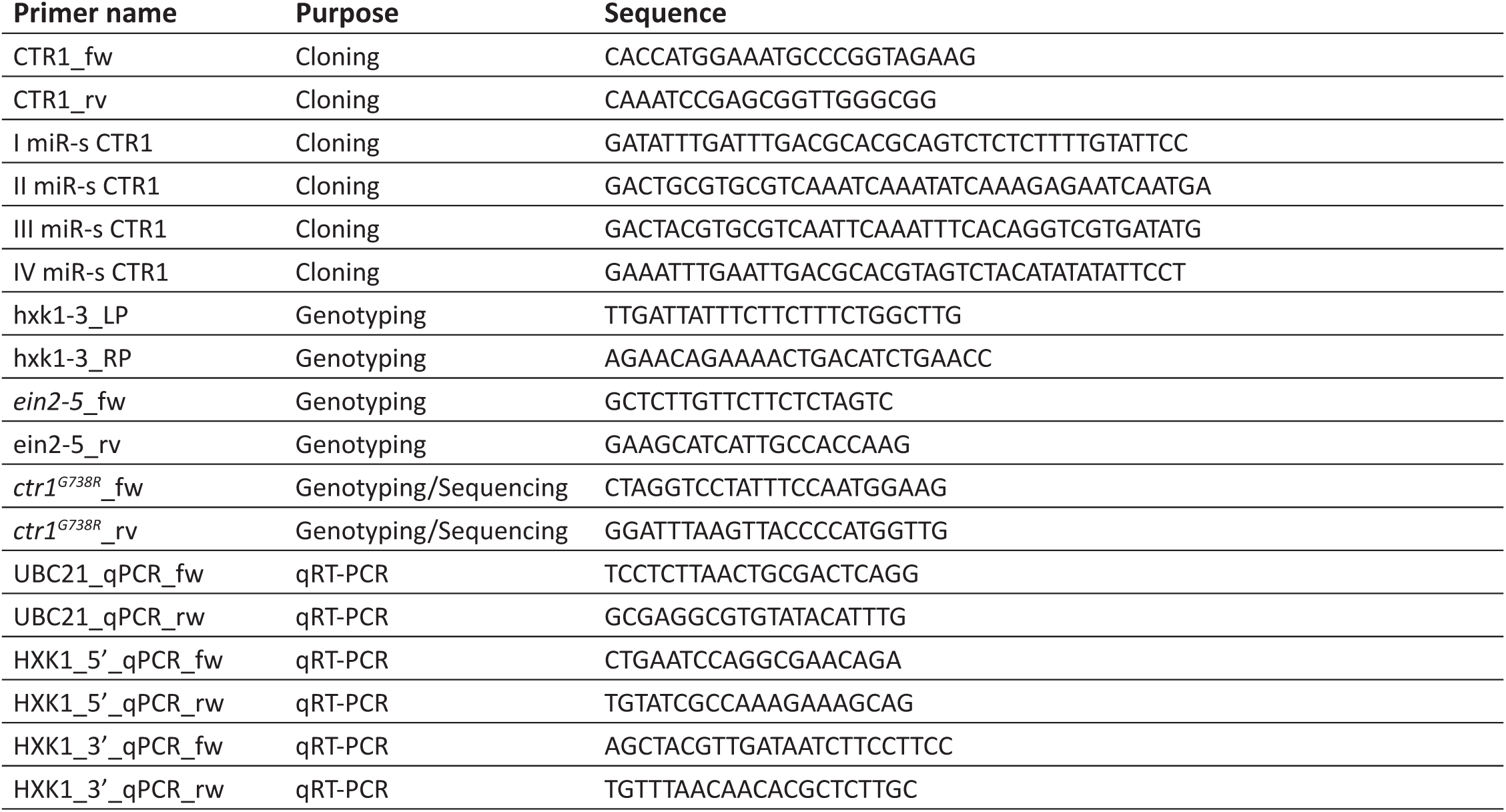

## ACKNOWLEDGMENTS

We thank Dr. Markus Grebe for providing the seeds of *ctr1-btk*, Dr. Wenrong He and Dr. Hongwei Guo for providing the seeds for *ein3eil1ebf1ebf2* and *EIN3ox*, Dr. Jiaying Zhu and Dr. Zhiyong Wong for providing the seeds of *gin2*, and Dr. Jose Alonso for providing the seeds of *ein2-5* and *ctr1-1*. We also thank Katelyn McKown, Gabriel Amador, and other members of the Bergmann lab for valuable feedback on the manuscript.

## ADDITIONAL INFORMATION

## Funding

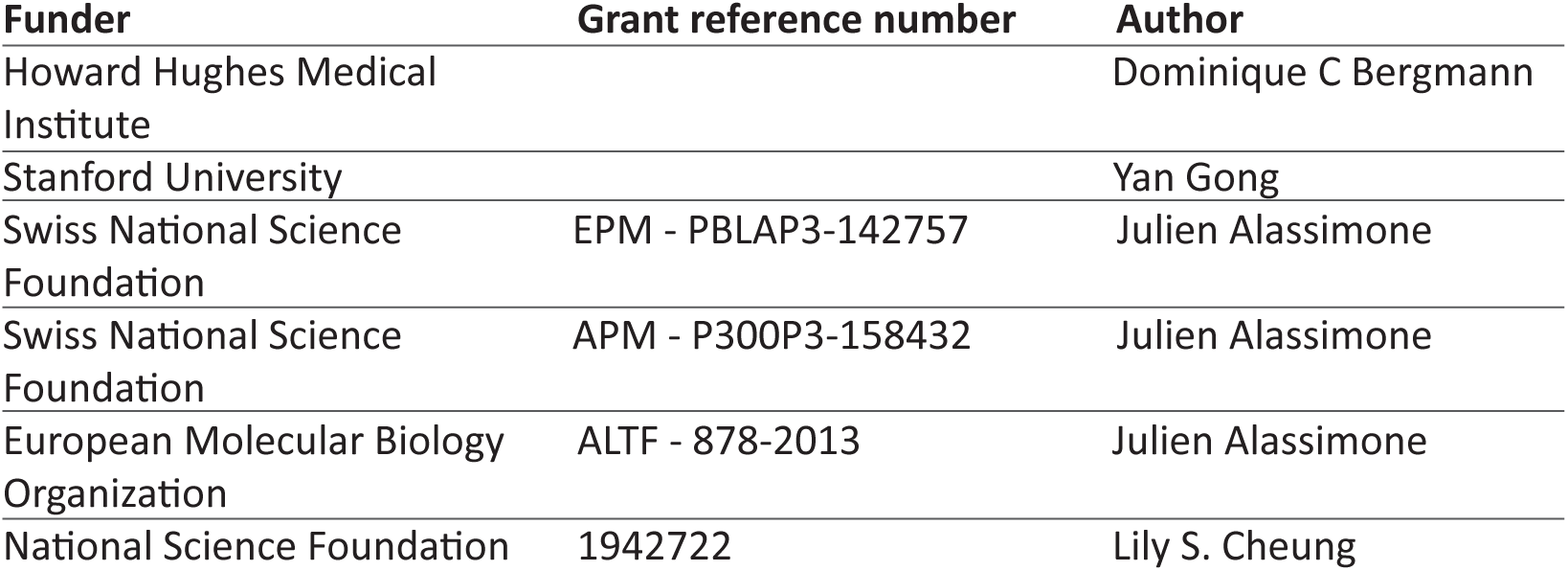

## Author contributions

Yan Gong, Conceptualization, Biological experiments and imaging, Data curation, Design of image quantification pipeline, Formal analysis, Image analysis, Interpretation of results, Methodology, Validation, Visualization, Writing—original draft; Julien Alassimone, Conceptualization, Biological experiments and imaging, Interpretation of results; Rachel Varnau, Biological experiments and imaging, Image analysis, Interpretation of results; Nidhi Sharma, Biological experiments and imaging, Interpretation of results; Lily S. Cheung, Design of image quantification pipeline, Interpretation of results, Methodology, Visualization, Dominique C. Bergmann, Conceptualization, Data curation, Formal analysis, Writing—original draft, Design of image quantification pipeline, Interpretation of results, Supervision, Funding acquisition, Project administration.

---

**Figure 1—figure supplement 1.**
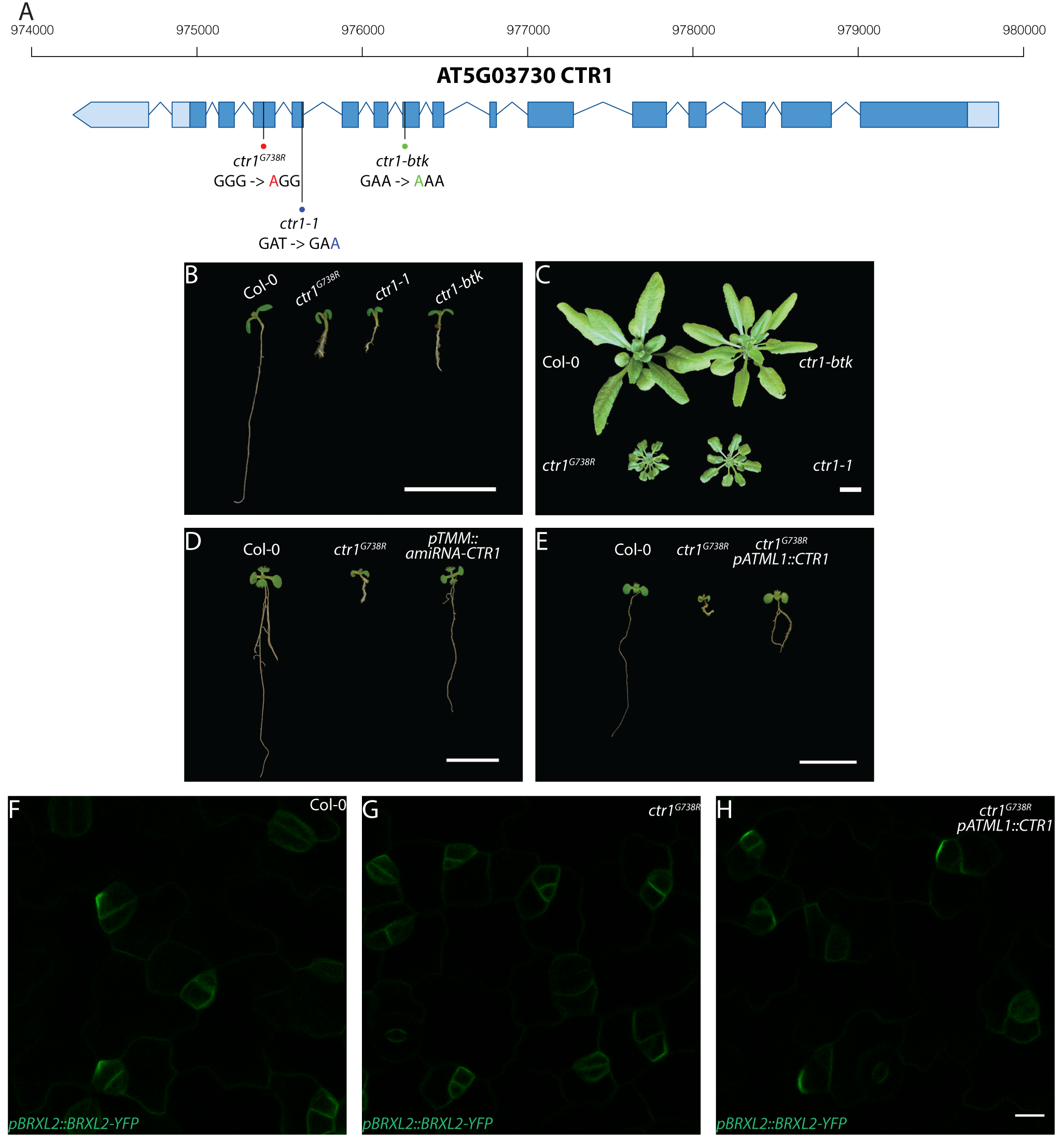
Molecular description of *ctr1*^*G738R*^ and other alleles, and whole plant phenotypes resulting from *ctr1* mutants, artificial microRNA knockdown, and shoot epidermal-only expression of *CTR1*. (A) Diagram of *CTR1* (At5g03730) locus and mutation sites for *ctr1*^*G738R*^, *ctr1-1*, and *ctr1-btk*. The *ctr1*^*G738R*^, *ctr1-1*, and *ctr1-btk* sites are marked by red, blue, and green dots respectively, and the corresponding nucleotide substitution site for each of mutations is labelled with the respective color. (B) Phenotypes of 7 dpg Col-0, *ctr1*^*G738R*^, *ctr1-1*, and *ctr1-btk* seedlings grown on ½ MS media without sugar (C) Phenotypes of 28 dpg Col-0, *ctr1*^*G738R*^, *ctr1-1*, and *ctr1-btk* plants grown on soil. (D) Phenotypes of 10 dpg Col-0, *ctr1*^*G738R*^, and *pTMM::amiRNA-CTR1* seedlings grown on ½ MS media without sugar. (E) Phenotypes of 7 dpg Col-0, *ctr1*^*G738R*^, and *ctr1*^*G738R*^ *pATML1::CTR1* rescue seedlings grown on ½ MS media without sugar. (F-H) Confocal images of *pBRXL2::BRXL2-YFP* (green) in 4 dpg cotyledons grown on ½ MS plates. (F) Col-0, (G) *ctr1*^*G738R*^, and (H) *ctr1*^*G738R*^ *pATML1::CTR1* rescue. Scale bar in B-E, 1 cm; H 10 µm.

**Figure 1—figure supplement 2.**
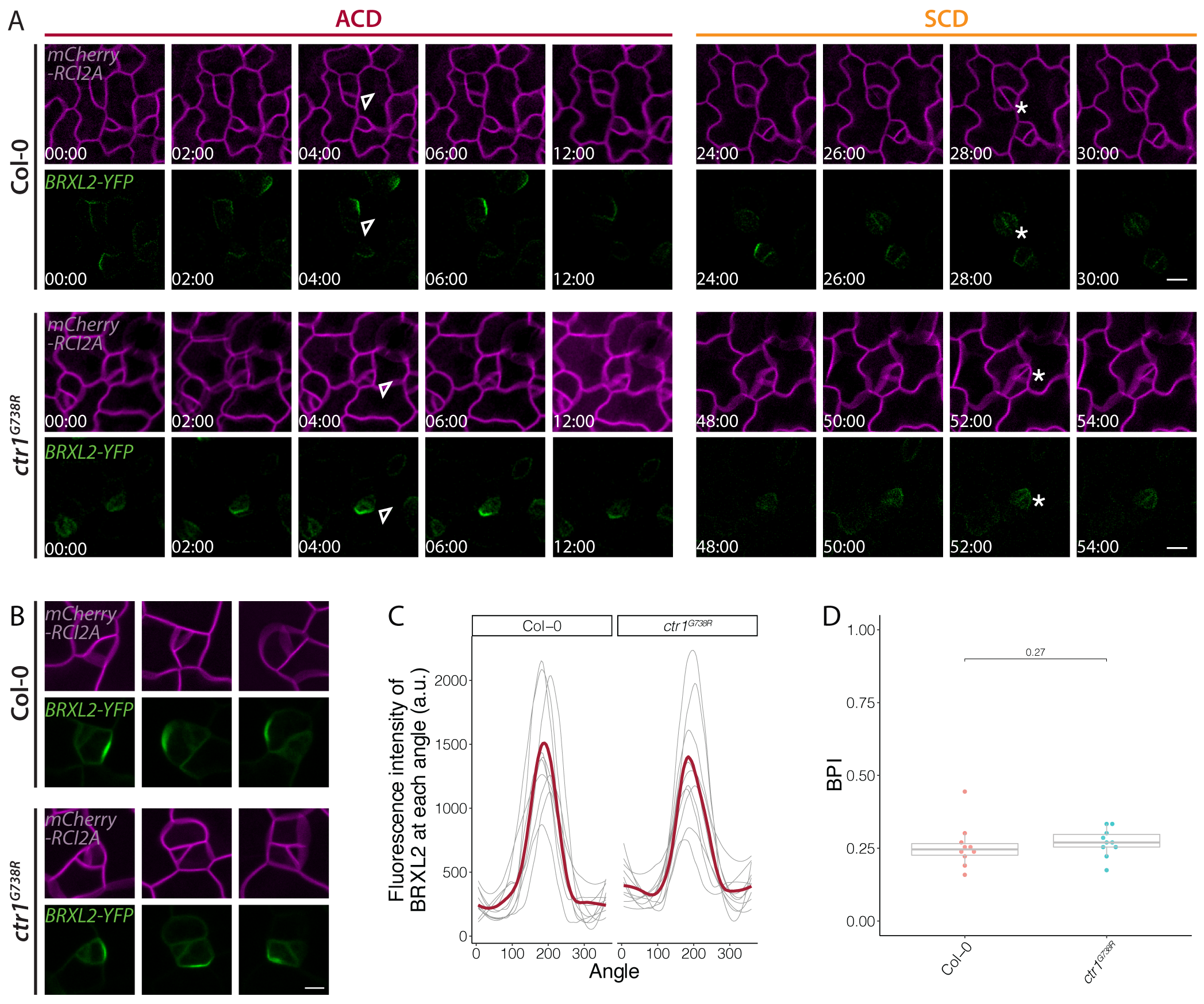
Time-lapse imaging of BRXL2 dynamics during stomatal lineage divisions and additional POME quantification of BRXL2 polarity in Col-0 and *ctr1*^*G738R*^. (A) Time-lapse confocal imaging of BRXL2 localization during an ACD (left) and a consecutive SCD (right) in Col-0 (top) and *ctr1*^*G738R*^ (bottom) cotyledons. *pBRXL2::BRXL2-YFP* and *pATML1::RCI2A-mCherry* are shown in green and magenta), and white numbers in frames indicate hours:minutes relative to first frame. Division types are marked by asterisks (GMC divisions) and triangles (amplifying divisions). (B-D) Cell polarity in BRXL2 polarized cells of 4 dpg Col-0 and *ctr1*^*G738R*^ cotyledons. Three example cells per each genotype are shown in (B). BRXL2 signal intensity distribution from quantified cells (C) and the BPI measurements of these cells (D) are shown. In (C), each thin line represents a single cell and the regression models for both genotypes are shown in cardinal red thick lines (n=10 cells/genotype in C-D). Scale bar in A-B,10 μm.

**Figure 2—figure supplement 1.**
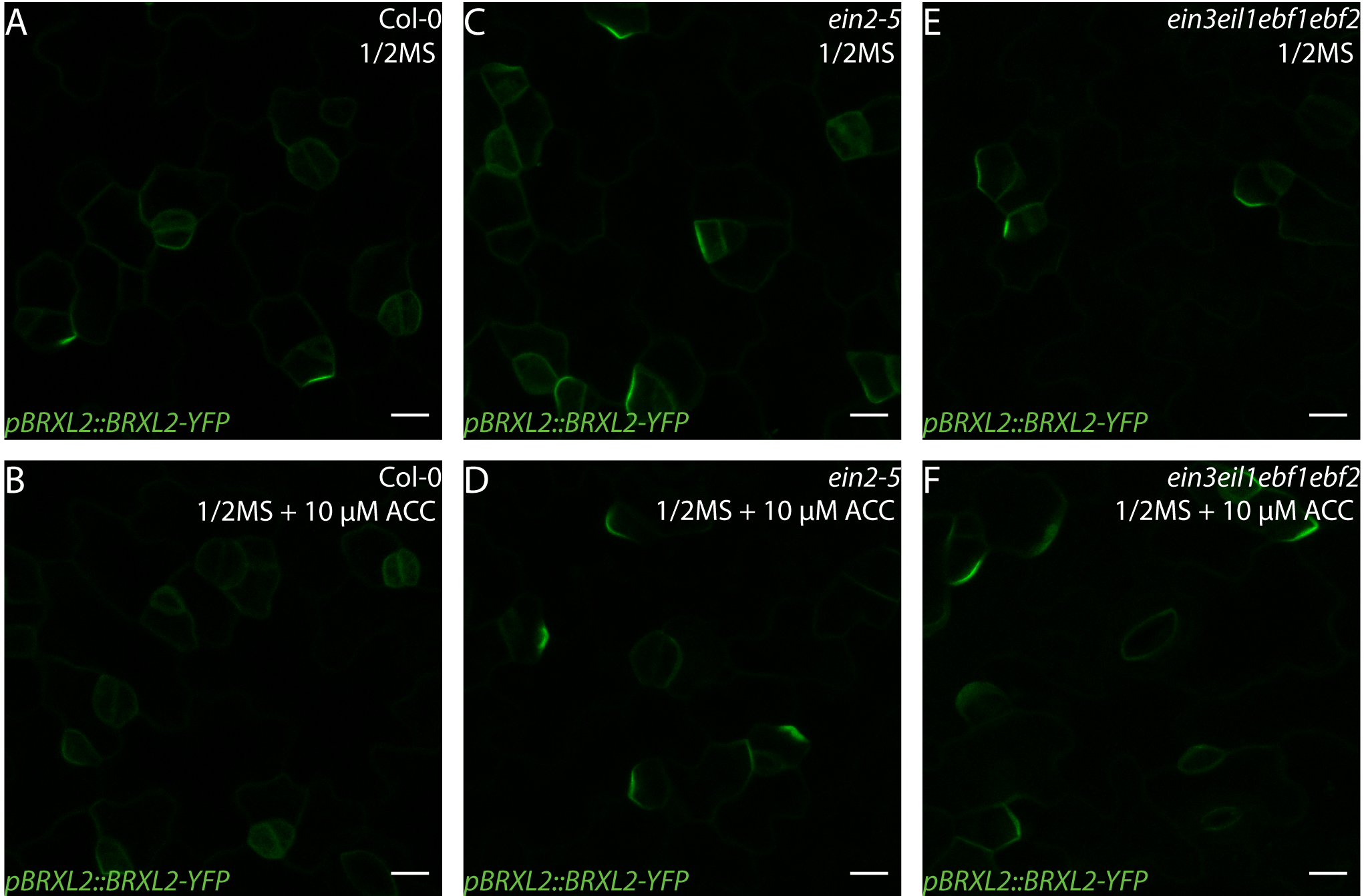
ACC does not affect BRXL2 polarity in ethylene insensitive mutants. (A-F) Confocal images of *pBRXL2::BRXL2-YFP* (green) in 4 dpg cotyledons grown on ½ MS plates with or without 10 µM ACC. (A-B) Col-0, (C-D) *ein2*, and (E-F) *ctr1-1*. (A, C, and E) ½ MS and (B, D, and F) ½ MS + 10 µM ACC. Scale bars in A-F, 10 µm.

**Figure 3—figure supplement 1.**
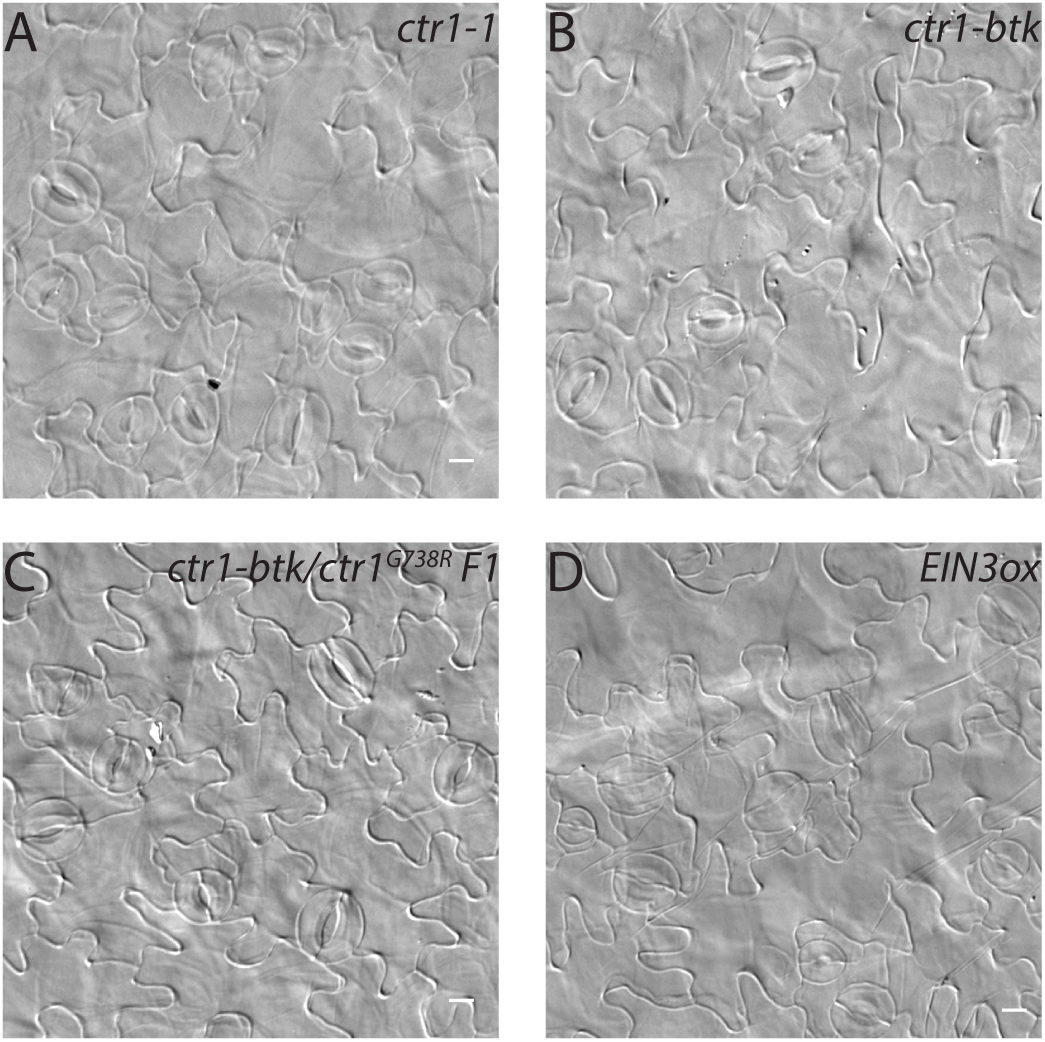
Additional phenotypes of cotyledon epidermis of ethylene signaling mutants. (A-D) DIC images of the epidermis in 14 dpg cotyledons of (A) *ctr1-1*, (B) *ctr1-btk*, (C) *ctr1-btk*/*ctr1*^*G738R*^ F1, and (D) *EIN3ox* grown on ½ MS media without sugar. Scale bars in A-D, 10 µm.

**Figure 4—figure supplement 1.**
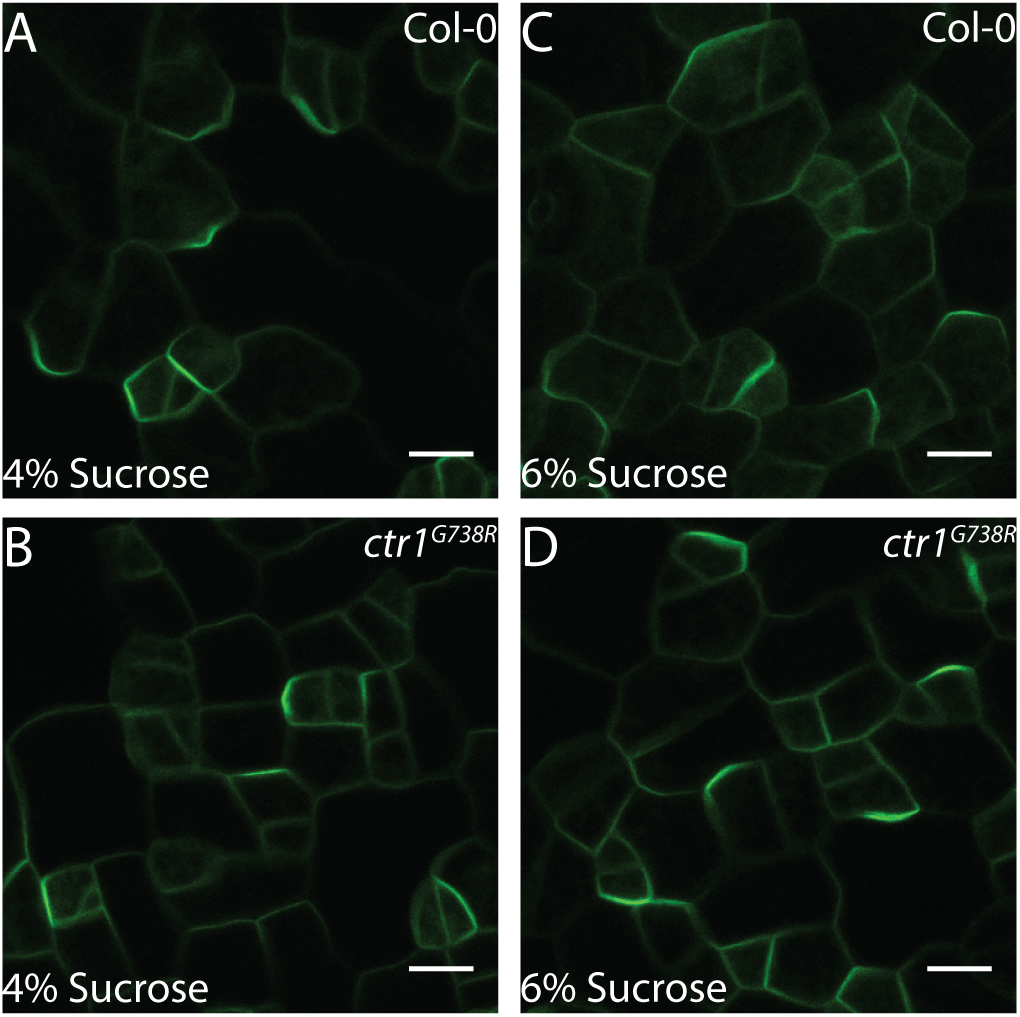
BRXL2 localization of Col-0 and *ctr1*^*G738R*^ grown on high levels of sucrose. (A-D) Confocal images of *pBRXL2::BRXL2-YFP* (green) in 4 dpg cotyledons grown on ½ MS plates with 4% or 6% sucrose. (A, C) Col-0 and (B, D) *ctr1*^*G738R*^. (A, B) ½ MS + 4% sucrose and (C, D) ½ MS + 6% sucrose. Scale bars in A-D, 10 µm.

**Figure 5—figure supplement 1.**
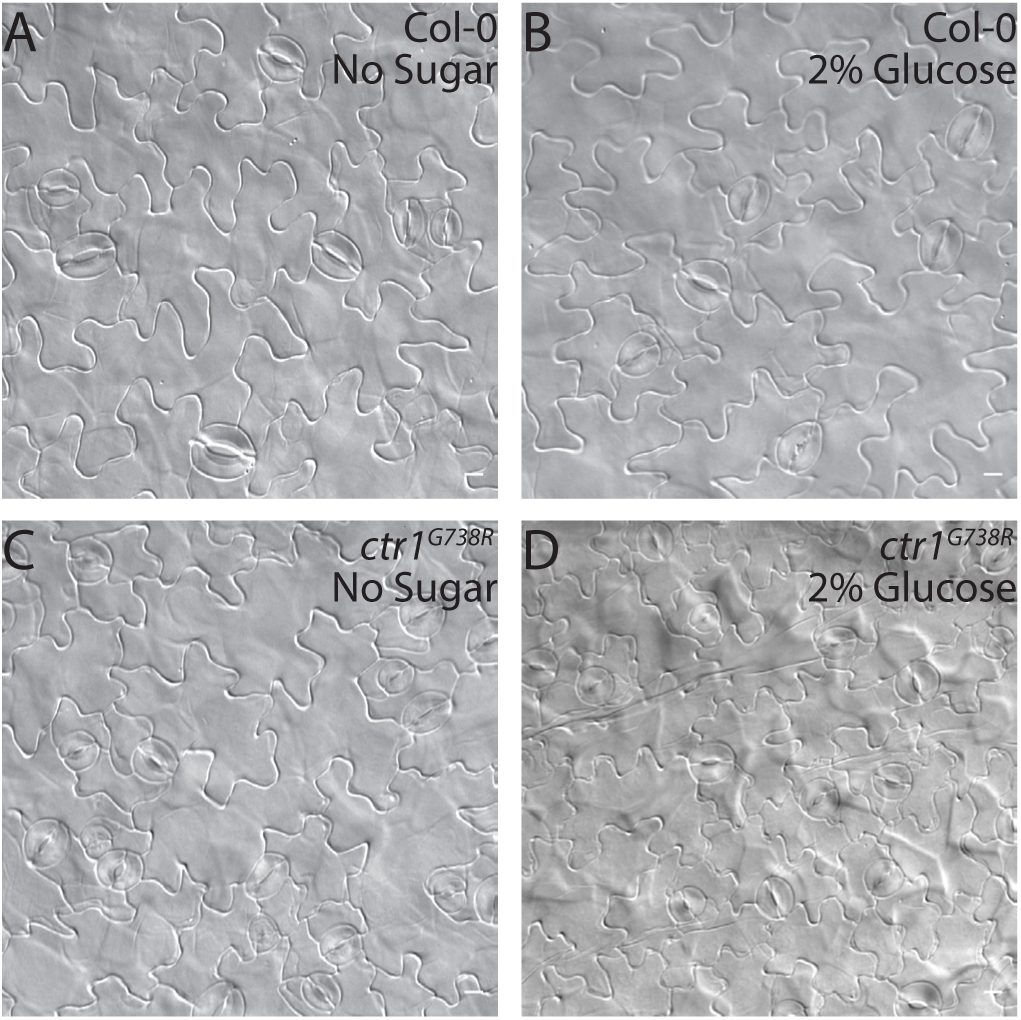
DIC images of cotyledons from seedlings grown on ½ MS media with or without 2% glucose. (A-D) DIC images of the epidermis in 14 dpg cotyledons grown on ½ MS media with or without 2% glucose. (A, B) Col-0 and (C, D) *ctr1*^*G738R*^. (A, C) ½ MS and (B, D) ½ MS + 2% glucose. Scale bars in A-D, 10 µm.

**Figure 5—figure supplement 2.**
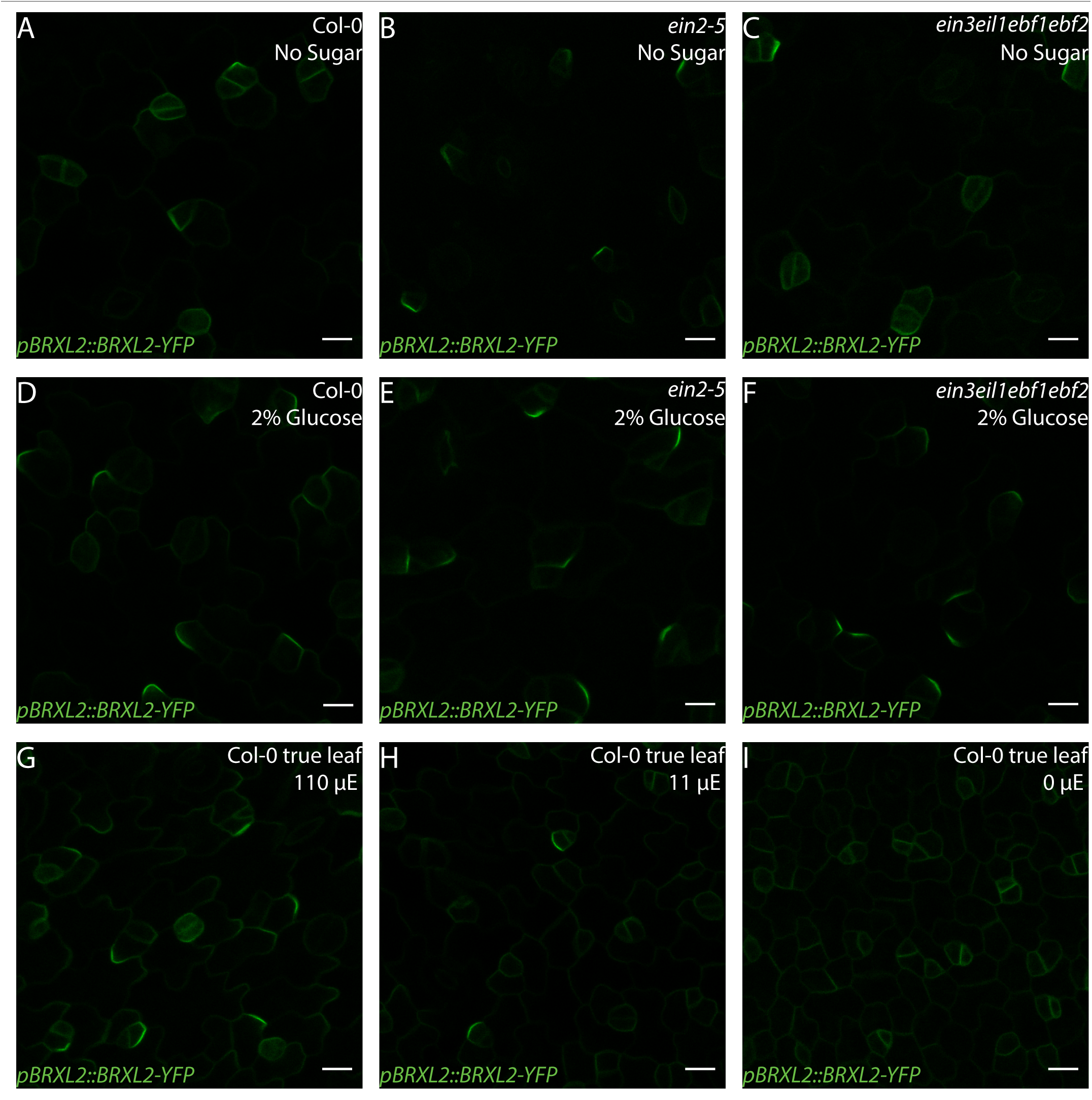
BRXL2 localization pattern in Col-0 and ethylene insensitive mutants under different light and sugar treatment regimes. (A-F) Confocal images of *pBRXL2::BRXL2-YFP* (green) in 4 dpg cotyledons grown on ½ MS plates with or without 2% glucose. (A, D) Col-0, (B, E) *ein2*, and (C, F) *ein3eil1ebf1ebf2*. (A-C) ½ MS and (D-F) ½ MS + 2% glucose. POME quantifications of BPI from A-F are shown in Figure 5C. (G-I) BRXL2 in 9 dpg Col-0 true leaves grown on ½ MS plates under different light condition. Col-0 seedlings are grown under 110 μE normal light condition for 7 days and then transferred to different light intensity conditions for 2 days. Light intensity in G-I: (G) 110 µE., (H) 10 µE, and (I) 0 µE. POME quantifications of BPI from G-I are shown in Figure 5F. Scale bars in A-I, 10 µm.

**Figure 5—figure supplement 3.**
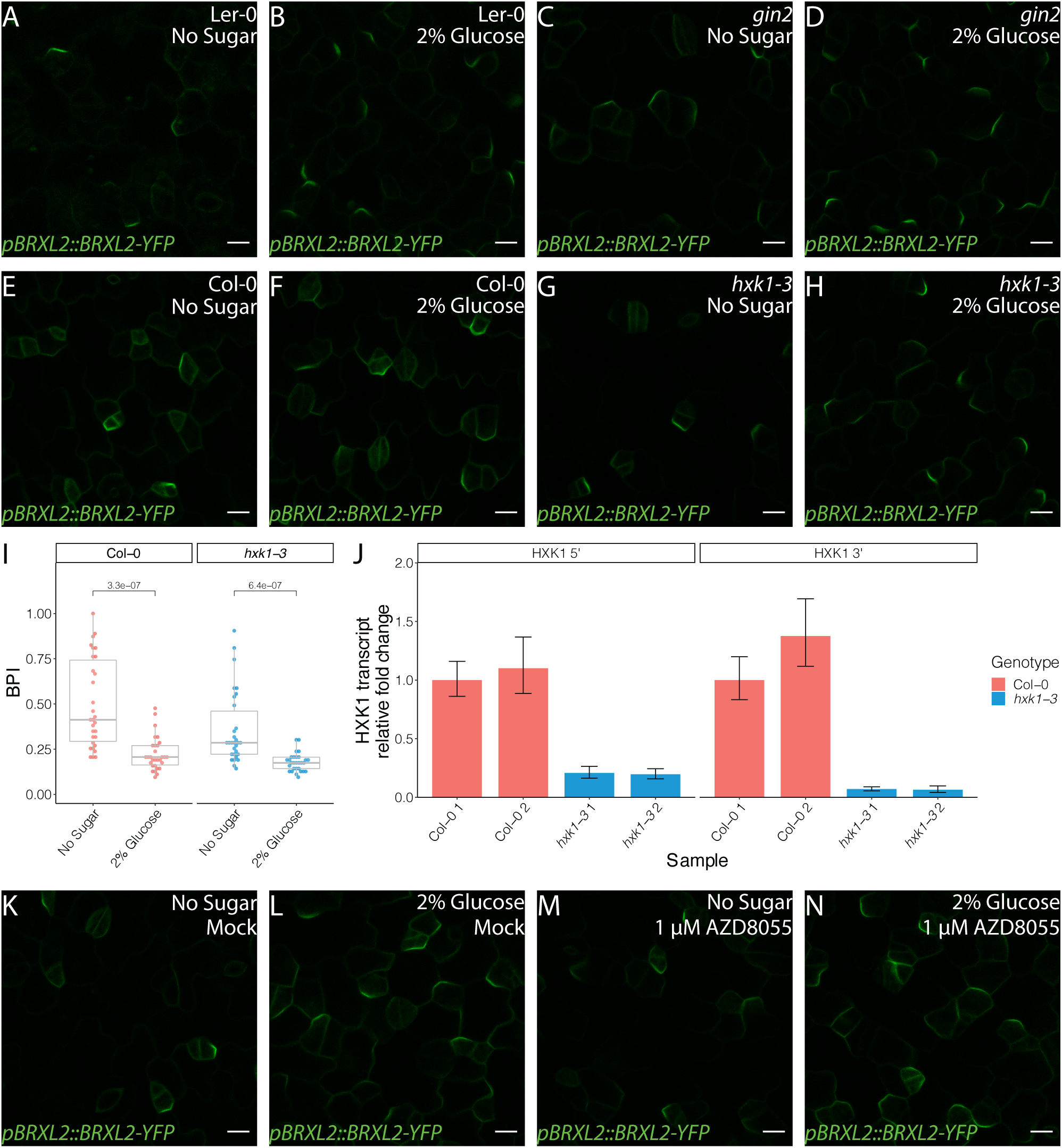
Glucose control of BRXL2 polarity is independent of HXK1 and TOR signaling. (A-H) Confocal images of *pBRXL2::BRXL2-YFP* (green) in 4 dpg cotyledons of different *hxk1* mutants and their corresponding wild-type controls grown on ½ MS plates with or without 2% glucose. (A, B) Ler-0, (C, D) *gin2*, (E, F) Col-0, and (G, H) *hxk1-3*. (A, C, E, and G) ½ MS and (B, D, F, and H) ½ MS + 2% glucose. POME quantifications of BPI from A-D are shown in Figure 5D. (I) POME quantifications of BPI from E-J conditions (n=30 cells/genotype) (J) qRT-PCR analysis of *HXK1* transcript level in 7 dpg Col-0 and *hxk1-3* seedlings. Relative transcript level of *HXK1* 5’ end (left plot) and 3’ end (right plot) are shown. Expression values are normalized to the control gene *UBC21* and are relative to the expression of first Col-0 replicate. (K-N) BRXL2 localization pattern in 4 dpg Col-0 treated with TOR inhibitor AZD8055 and/or 2% glucose. (K, M) ½ MS and (L, N) ½ MS + 2% glucose. (K, L) mock treatment and (L, N) 1µM AZD8055 treatment. POME quantifications of BPI from K-N are shown in Figure 5E. All p-values are calculated by Mann-Whitney test. Scale bars in A-H and K-N, 10 µm.

**Figure 6—figure supplement 1.**
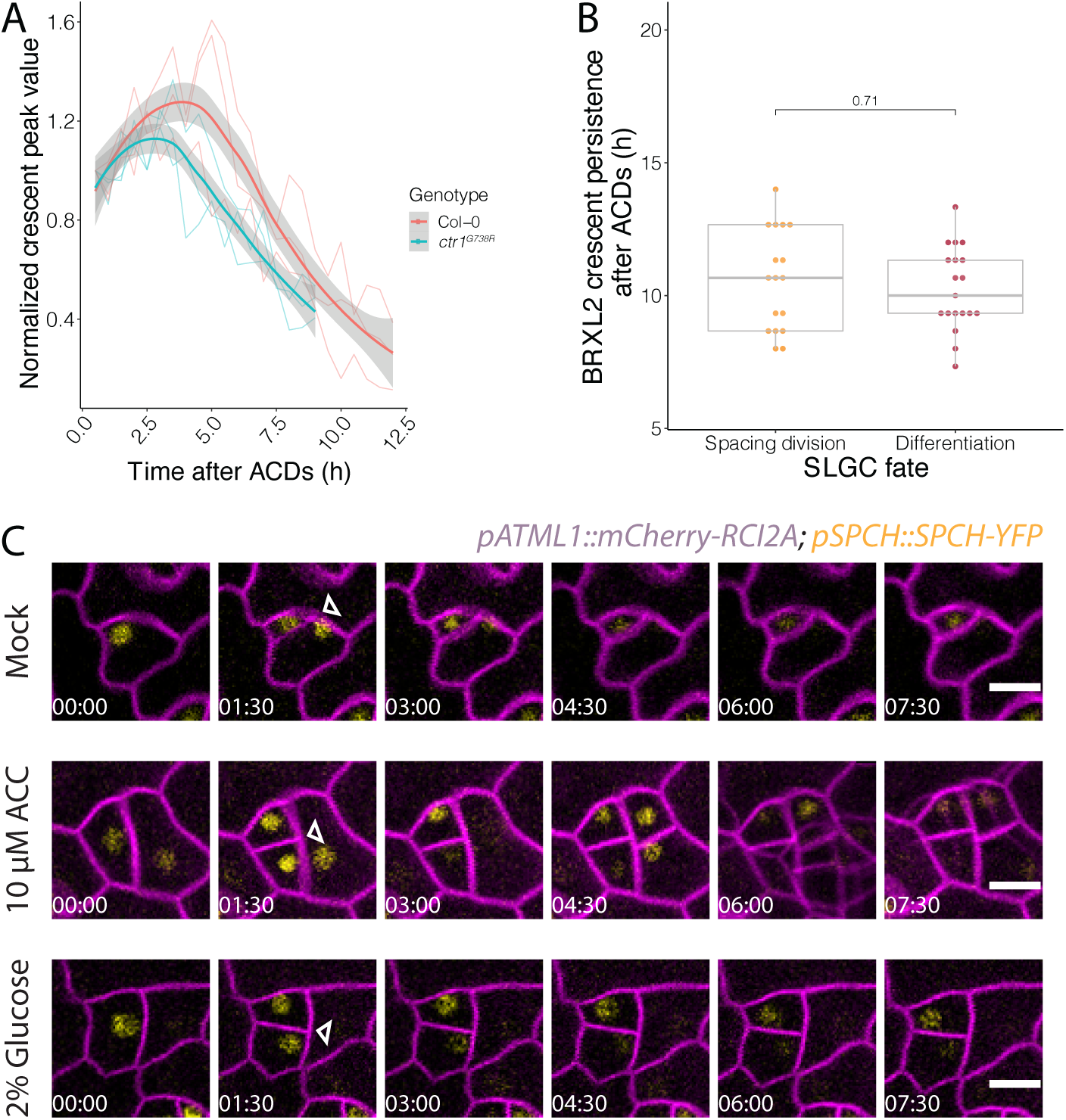
Additional characterizations of BRXL2 and SPCH dynamics during ACDs. (A) Normalized peak value of BRXL2 crescents measured by POME over time from cells in Col-0 and *ctr1*^*G738R*^ (n=3). Individual measurements per cell are shown in thin lines and the respective trend per each genotype with 0.95 confidence interval is indicated as the thick line with gray band. BPI measurements of the same cells are shown in Figure 6B. (B) Persistence of post-ACD BRXL2 in SLGCs grouped based on whether they undergo spacing divisions or differentiation (no spacing division). (C) Time-lapse images of SPCH dynamics in 3 dpg mock, 10 µM ACC, or 2% glucose treated Col-0 seedlings. *pSPCH::SPCH-YFP* and *pATML1::RCI2A-mCherry* are shown in yellow and magenta. Each cell division is marked by a triangle, and white numbers in frames indicate hours:minutes relative to first frame. p-values are calculated by Mann-Whitney tests. Scale bars in C, 10 µm.

